# The mycobacterial nucleoid-associated protein NapM exhibits stress-induced septal localization and modulates cell envelope gene expression

**DOI:** 10.1101/2024.11.29.625984

**Authors:** Kornel Milcarz, Joanna Hołówka, Jakub Pawełczyk, Mariola Paściak, Yaroslav Lavrynchuk, Przemysław Płociński, Jolanta Zakrzewska-Czerwińska

**Affiliations:** Department of Molecular Microbiology, University of Wrocław, Wrocław, Poland; Laboratory of Genetics and Physiology of Mycobacterium, Institute of Medical Biology, Polish Academy of Sciences, Łódź, Poland; Department of Immunology of Infectious Diseases, Hirszfeld Institute of Immunology and Experimental Therapy, Polish Academy of Sciences, Wrocław, Poland; Department of Immunology and Infectious Biology, Faculty of Biology and Environmental Protection, University of Łódź, Łodź, Poland

## Abstract

The bacterial chromosome is organized in a hierarchical and dynamic manner to facilitate various DNA transactions. Nucleoid-associated proteins (NAPs) are the most abundant small-scale chromosomal organizers, playing roles in maintaining chromosomal DNA integrity, gene regulation, DNA replication, and DNA repair. In this study, we characterize the recently identified mycobacterial NAP, NapM in *Mycobacterium smegmatis*. Our study shows that NapM exhibits a distinctive septal localization in response to stress affecting cell envelope integrity and modulates the expression of approximately one-third of *M. smegmatis* genes, including those involved in cell envelope biosynthesis. Our findings suggest that NapM regulates mycobacterial cell division under stress, enabling this saprophyte to adapt to constantly changing environmental conditions.

## Introduction

Emerging evidence shows that the bacterial chromosome resembles eukaryotic chromatin having a highly organized and hierarchical structure (1). Although the chromosome must be tightly condensed to fit into the bacterial cell, certain regions need to be accessible to the protein machineries involved in basic DNA transactions, such as replication, transcription, and DNA repair. Different groups of proteins contribute to maintaining the hierarchical yet dynamic nucleoid structure. Among them are topoisomerases, which modulate chromosome topology (2); condensins, which contribute to long DNA fragment organization (3); and small basic proteins called nucleoid-associated proteins (NAPs) (1). NAPs organize the chromosome more locally by introducing bends (e.g., HU and IHF homologs), bridging DNA (e.g., H-NS, Fis), bunching and wrapping (e.g., IHF, H-NS), and stiffening (e.g., HU, Fis) (4). Moreover, they can bind DNA to act as global regulators of transcription (5–7).

While some NAPs exhibit structural and/or functional homology across various bacterial species, others are conserved within a single genus or species. The genus *Mycobacterium* possesses a distinct set of NAPs with unique functions, often beyond shaping chromosome architecture (e.g., regulation of chromosome replication and gene transcription) (8–10). Mycobacteria elongate apically and divide asymmetrically, such that chromosome is positioned closer to the new pole (8). Moreover, the structure of the mycobacterial chromosome is unusual within the bacterial world, as it has bead-like chromosomal domains, the formation of which remains unknown (11). The most abundant mycobacterial NAP, HupB (an HU homolog), presumably participates in pre-replication complex formation (12) and inhibits RecA-promoted strand exchange (13). The functional homolog of H-NS, Lsr2, together with its ortholog, MSMEG_1060, helps the cell adapt to unfavorable conditions, potentially by regulating gene transcription (e.g., genes involved in the synthesis of lipooligosaccharides; LOS) (10, 14). Additionally, Lsr2 promotes the exchange of replicative DNA polymerase during chromosomal DNA synthesis in *Mycobacterium smegmatis*, balancing mutagenesis and survival under DNA-damaging conditions, and contributing to the emergence of antibiotic (e.g., rifampicin) resistance (15). Mycobacterial IHF is essential for the integration of mycobacteriophage L5 (16), and its depletion results in chromosome shrinkage and replication inhibition (17). Additionally, certain NAPs are encoded only within the genomes of pathogenic mycobacteria; these include the virulence regulators, EspR (18), NapA, (19), and MDP2, which interacts with the polar growth determinant, DivIVA (20).

The recently identified nucleoid-associated protein, NapM, is a well conserved protein found in both saprophytic and pathogenic species of *Mycobacterium* (21). *M. smegmatis* NapM was shown to be involved in altering resistance to ethambutol (EMB) and rifampicin (antibiotics used to treat tuberculosis) (21). Interestingly, in *M. tuberculosis*, NapM inhibits DNA replication, presumably by binding the replication initiation protein, DnaA (22). Thus, NapM may facilitate the persistence of *M. tuberculosis* as non-replicating cells within macrophages during latent tuberculosis infections (21, 22). Although *M. smegmatis* and *M. tuberculosis* belong to the same genus, their occupied niches differ significantly. *M. smegmatis* primarily inhabits water and soil environments, often living symbiotically with other organisms, such as free-living amoebae (23). In immunocompromised patients, however, *M. smegmatis* can cause opportunistic infections that mainly affect the skin and soft tissues (24). It is unknown whether the NapM in saprophytic *M. smegmatis* is also involved in chromosome replication as in *M. tuberculosis*. Here, we show that in *M. smegmatis*, NapM modulates cell cycle dynamics and undergoes stress-induced relocalization to the septum, a phenomenon not previously observed in other NAPs. Moreover, we demonstrate that NapM functions as a pleiotropic transcriptional regulator that impacts over one-third of *M. smegmatis* genes, particularly those involved in cell envelope biosynthesis, leading to the accumulation of one of its components – phosphatidylinositol dimannoside (PIM_2_). These findings indicate that NapM exhibits a broader range of functions than its originally proposed role as a nucleoid-associated protein, raising questions about its classification within this group.

## Results

NapM is annotated as a member of the PadR family, a large group of transcriptional regulators that function as environmental sensors. PadR family members, including NapM possess two domains: an N-terminal DNA-binding winged helix-turn-helix (wHTH) domain and a C-terminal domain responsible for homodimerization (25). Although NapM occurs exclusively in *Mycobacterium* and is highly conserved within this genus (average identity – 92%, **Fig. S1A**), the phylogenetic tree of NapM proteins (**Fig. S1B**) suggests that there is some diversity in the C-terminal domain sequences between saprophytes (e.g., *M. smegmatis*) and pathogens (e.g., *M. tuberculosis*). Previous atomic force microscopy experiments suggested that NapM is a DNA-bridging protein that forms aggregates with looped dsDNA (21, 26). However, the literature lacked direct evidence that NapM can dimerize in *M. smegmatis*. To address this, we carried out Bacterial Two-Hybrid (BTH) experiments and found that NapM forms homodimers (**Fig. 1A**), and the dimer structure is presumably stabilized by interactions of the C-terminal domains, as further supported by AlphaFold predictions (**Fig. 1B**). Additionally, the recently uploaded crystal structure of NapM from *M. tuberculosis,* which shows the formation of homodimers, further corroborates our results (https://swissmodel.expasy.org/repository/uniprot/P71704).

**Figure 1.**
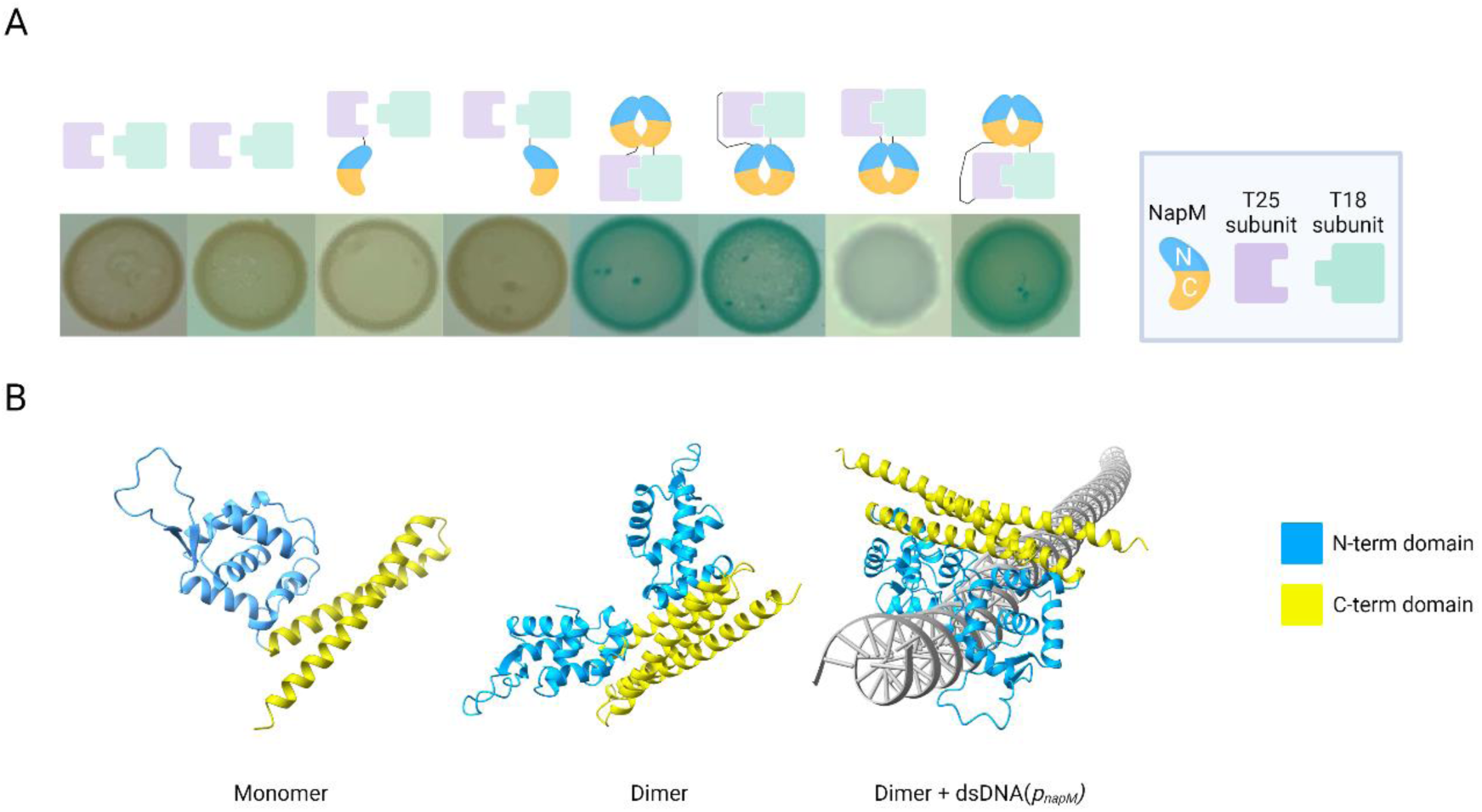
NapM forms homodimers. **A.** Bacterial Two-Hybrid (BTH) assay (spots) of the *M. smegmatis* NapM protein. **B.** AlphaFold (https://alphafoldserver.com/welcome) predictions of *M. smegmatis* NapM tertiary structure.

### NapM regulates cell cycle dynamics

To elucidate the function of NapM during *M. smegmatis* growth, we analyzed the single-cell morphology, chromosome organization, and membrane integrity of the *napM* deletion mutant (Δ*napM*) and a strain overproducing NapM protein (NapM↑) under optimal conditions. There was no apparent difference in the membrane continuity of the tested strains (**Fig. S2A**). However, the Δ*napM* cells were longer than the wild-type cells (4.4 ± 2.1 µm and 3.5 ± 1.4 µm, respectively; t(598) = 8.43, *p* = 2 × 10⁻¹⁶, n = 300 per group; see **Fig. 2A**), whereas the NapM↑ cells were shorter than Control↑ (wild-type strain with an empty pMV_pAMI_ vector after induction with 1% acetamide) cells (2.7 ± 0.7 µm and 3.3 ± 0.9 µm, respectively; t(598) = −9.11, *p* = 3.2 × 10⁻¹⁶, n = 300 per group; **see Fig. 2A**). The average cell width was similar across the strains (**Fig. 2B**), and no significant between-strain difference was observed in the asymmetry ratio of daughter cells (**Fig. 2C**). These data suggest that NapM may influence the cell elongation and/or cell envelope composition of *M. smegmatis* potentially through transcriptional regulation. Our experiments also revealed that changes in the NapM level slightly affected chromosome condensation, calculated as a percentage of cell length (76% ± 6 and 79% ± 5 for the deletion mutant and wild-type strain, respectively; t(98) = −2.72, *p* = 4.2 × 10⁻², n = 50 per group; **see Fig. S2B**). However, the nucleoid was longer in the Δ*napM* cells than in the wild-type cells (2.91 ± 0.88 µm and 3.68 ± 1.10 µm; t(198) = −5.47, p = 6.4 × 10⁻⁷, n = 100 per group; **see Fig. 2D**). In contrast, these parameters were not altered in NapM↑ cells compared to the Control↑ strain (**Fig. 2D, Fig. S2B**).

**Figure 2.**
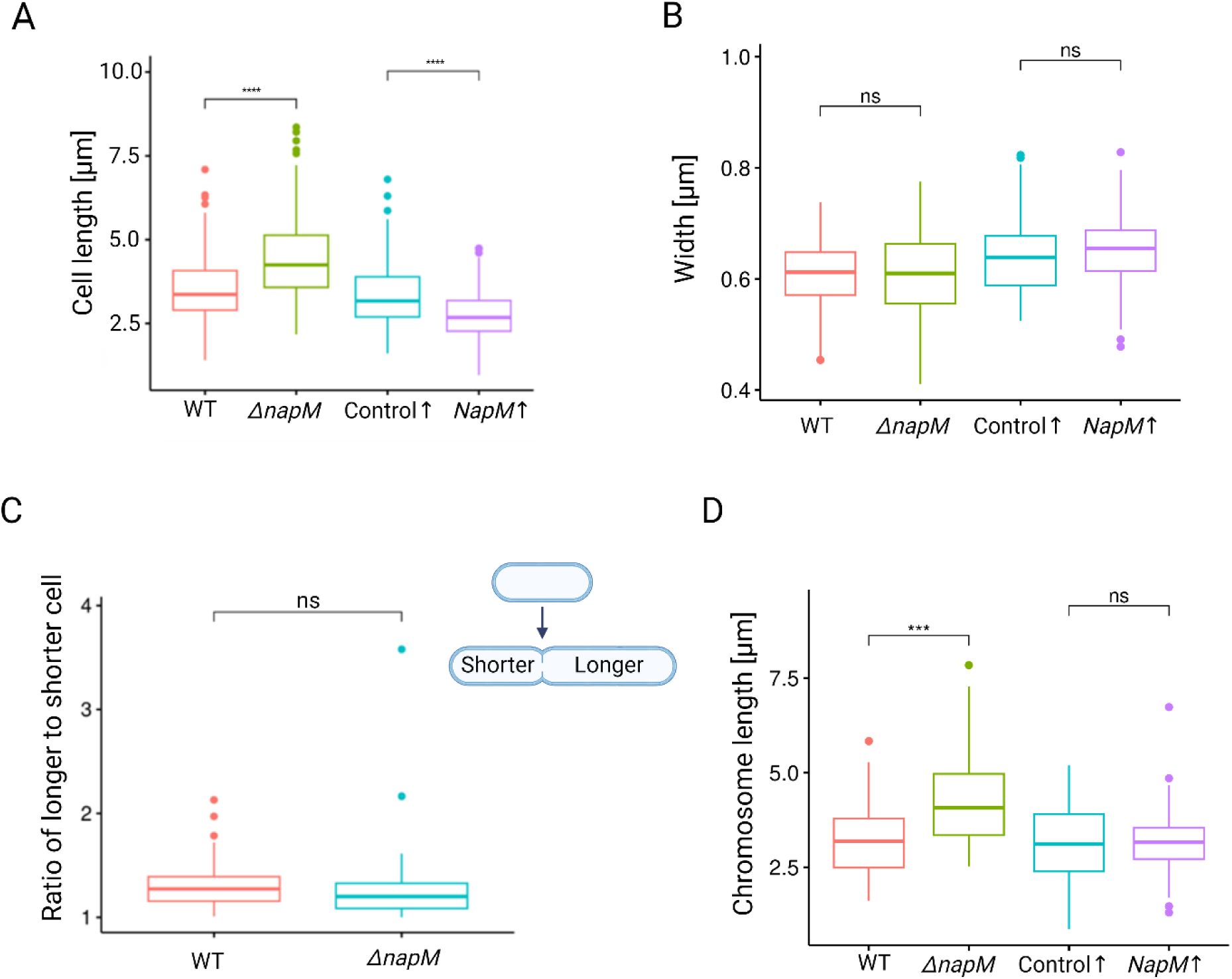
*napM* deletion alters the cell and chromosome length. Boxplots presenting the average cell lengths (**A**), cell width (**B**), and daughter cells asymmetry (**C**) for tested strains. **D.** Boxplots showing length of the chromosomes stained with Hoechst 33342 dye in analyzed strains.

To further investigate the role of NapM in cell cycle dynamics in *M. smegmatis*, we used the previously described chromosome replication marker, DnaN-mCherry (27) in the Δ*napM* and NapM↑ strains. Time Lapse Fluorescence Microscopy (TLFM) experiments revealed that, under optimal conditions, Δ*napM* cells exhibited a slight delay in replication time (C phase) relative to wild-type cells (125.58 ± 21.30 min and 120.20 ± 16.84 min, respectively; t(608) = 3.46, p = 5.4 × 10⁻⁴, n = 305 per group; see **Fig. S3A**) but no significant difference was observed in the time from replication termination to the initiation of the new replication round in daughter cells (BD phase) (**Fig. S3B**). Interestingly, under nutrient-restriction conditions (limited nutrients availability, see M&M), Δ*napM* exhibited delays in both the C phase (131.99 ± 18.42 min vs. 124.34 ± 1.42 min; t(198) = 4.14, p = 3.6 × 10⁻⁶, n = 272 for Δ*napM*, 267 for WT) and, particularly, the BD phase compared to the WT strain (57.94 ± 26.94 min vs. 45.73 ± 19.54 min; t(198) = 3.67, p = 1 × 10⁻¹⁰ n = 326 for Δ*napM*, and 314 for WT, see **Fig. 3**). Moreover, doubling time for Δ*napM* was extended in comparison to WT (190.29 ± 34.69 min and 170.67 ± 28.27 min, respectively; t(498) = 6.93, p = 2.2 × 10⁻¹², n = 250).

**Figure 3.**
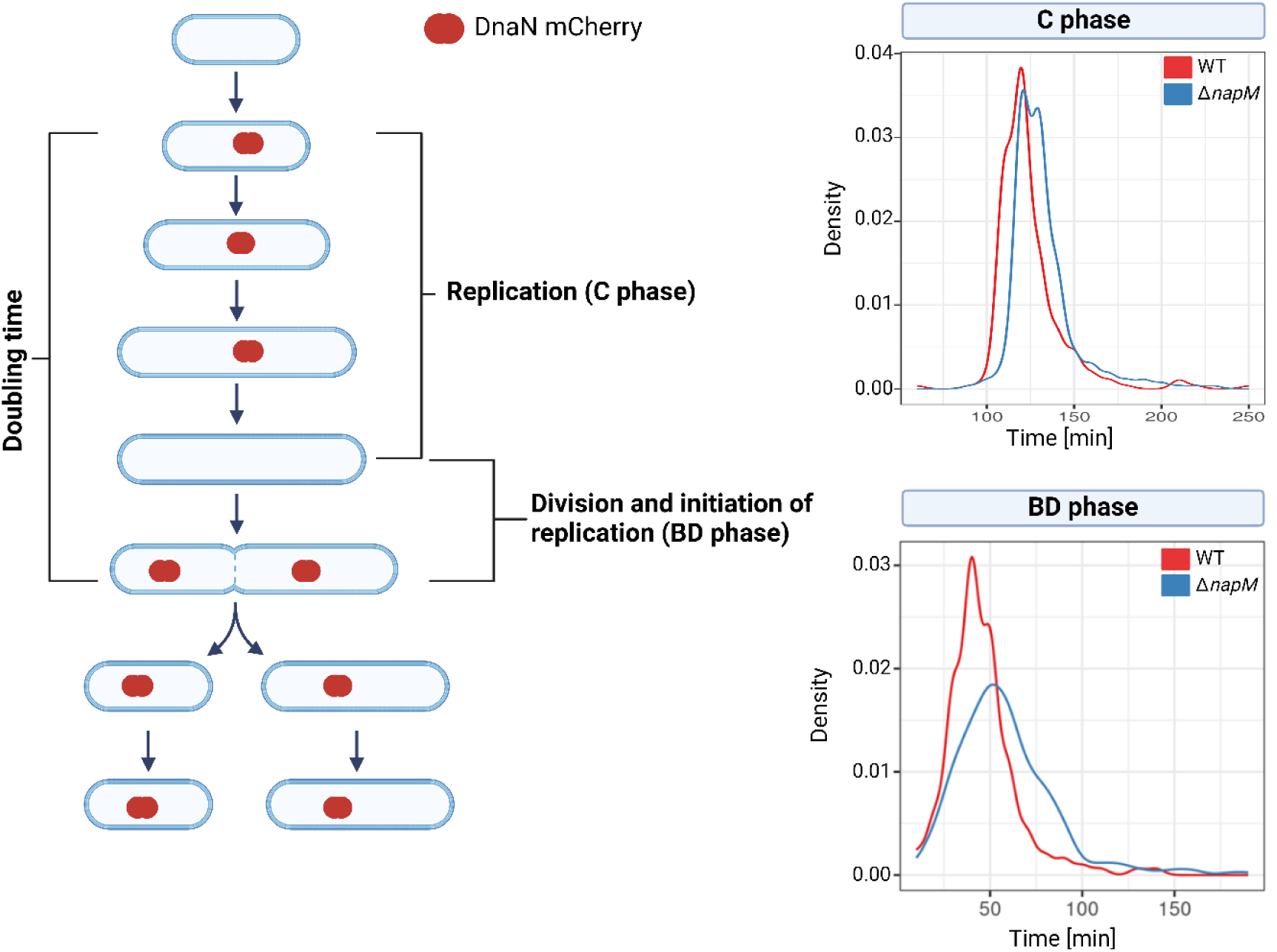
Replication dynamics is altered in the *ΔnapM* strain under nutrient-restricted conditions. The left panel shows a schematic representation of the phases of mycobacterial cell cycle. The right panel displays a density plot of the durations of the C and BD phases in the wild-type (WT) and *ΔnapM* strains.

Together, these results suggest that NapM influences cell division dynamics in *M. smegmatis* and its lack slow down replication process under nutrient restriction. This indicates that NapM plays a crucial role in regulating the cell cycle in response to nutrient depletion in the saprophytic *M. smegmatis*.

### NapM acts as a pleiotropic transcriptional regulator

Since NapM belongs to the PadR family of transcription factors, we investigated whether NapM regulates gene transcription. To this end, we performed RNA-Seq (GEO Accession GSE275852) to compare the global transcription profiles of Δ*napM* and the wild-type strain in optimal conditions. The results revealed that deletion of *napM* significantly altered the expression of over 2,400 genes (with analysis threshold set to abs log_2_FC = 1, FDR = 0.05) in exponentially growing *M. smegmatis* cells. Specifically, 1609 genes exhibited increased expression, while 816 showed decreased expression when *napM* was deleted, indicating that NapM may primarily function as a transcriptional repressor (**Fig. 4A**). Although NapM-regulated genes were distributed across the entire chromosome, certain chromosomal regions were particularly affected, with the most substantial changes occurring around the *oriC* region suggesting its potential role in regulation of housekeeping genes. Additionally, we observed moderate changes along the transcripts of genes allocated within chromosomal arms, and the least changes near the *ter* region (**Fig. 4B**). The transcriptional alterations in the Δ*napM* strain were visualized using a volcano plot (**Fig. 4C**), highlighting the most significant changes in transcripts such as *panB, glpA/glpD, MSMEG_6760, glpK, ino1, L-lactate 2-monooxygenase, MSMEG_4208, MSMEG_4209, MSMEG_4211,* and *MSMEG_4207.* Using the KEGG database, we classified the genes with altered transcript levels from the RNA-Seq data (**Fig. 4D**). The most significant changes compared to the wild-type strain were associated with processes such as mismatch repair, homologous recombination, base excision repair, and DNA replication (*dnaN, dnaQ*) (**Fig. 4D**). Of particular interest were gene groups potentially linked to the observed elongated cell phenotype, including those involved in cell division (*wag31, crgA, MSMEG_6171*) and cell envelope synthesis (*MSMEG_2934, MSMEG_0359, MSMEG_3859*) (**Table S4**).

**Figure 4.**
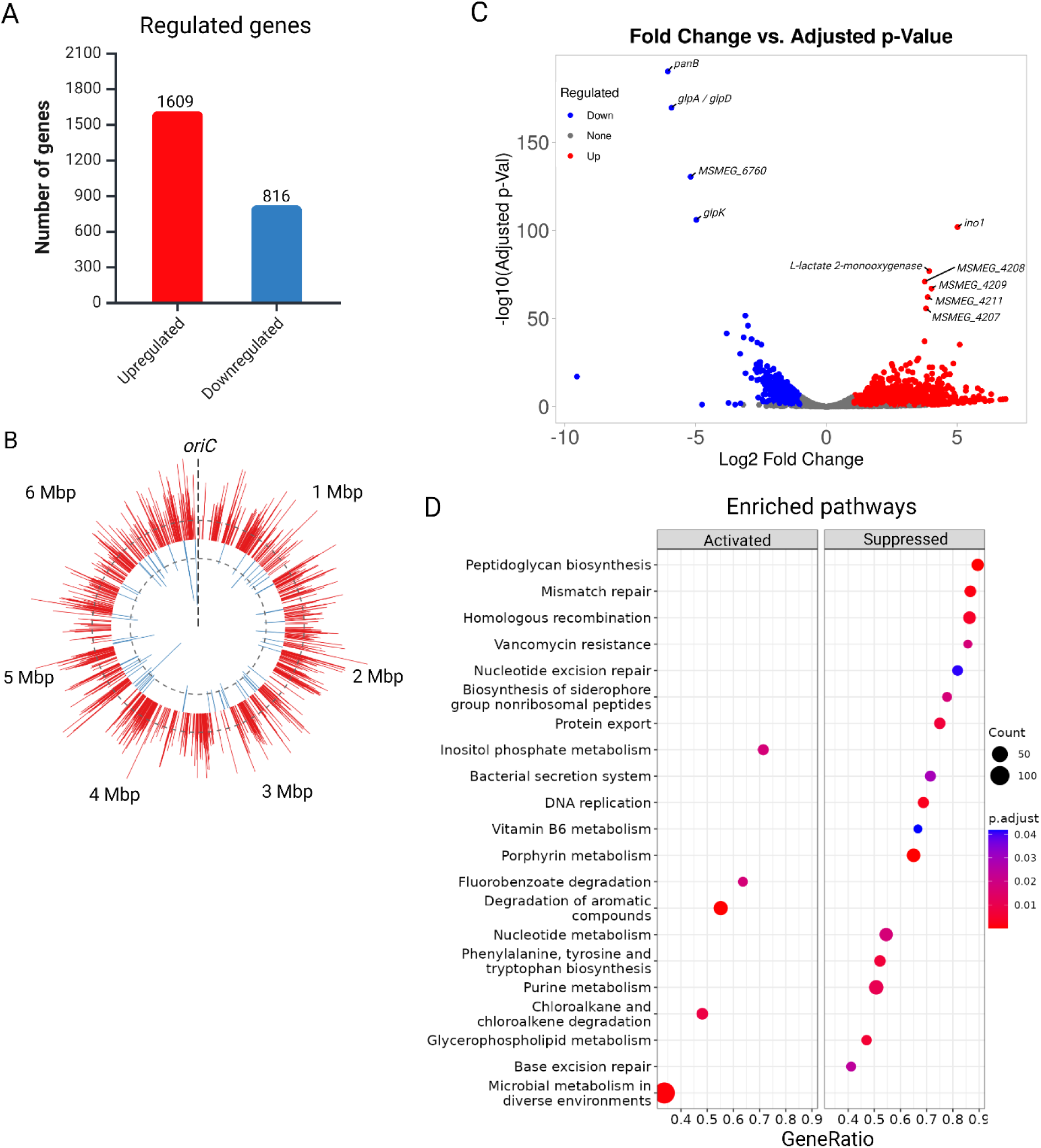
*napM* deletion alters the expression of genes across the chromosome. **A.** Number of genes regulated in response to *napM* deletion. **B.** Chart presenting transcriptomic changes relative to gene position in the genome of the *ΔnapM* strain. **C.** Enriched pathways in the *ΔnapM* strain (KEGG database). **D.** Volcano plot for the *ΔnapM* strain with highest-scoring genes annotated.

Given our observation that NapM may influence cell elongation and division (**Fig. 2A**), we further focused our analysis on transcriptomic changes related to cell envelope biosynthesis and cell division. Notably, NapM functions as an activator for cell envelope genes; in the Δ*napM* strain, peptidoglycan (PG) biosynthesis genes were downregulated across all synthesis steps, including cytoplasmic precursor synthesis, the formation and subsequent translocation of lipid-linked intermediates across the cytoplasmic membrane, and PG polymerization and cross-linking (**Table S4**). Additionally, several genes involved in phosphatidylinositol-containing polar lipid biosynthesis, spanning multiple pathways starting from phosphatidylinositol (PI) synthesis, through phosphatidylinositol mannosides (PIMs) formation, to their glycosylation into lipomannan (LM) and lipoarabinomannan (LAM), exhibited downregulated expression in *ΔnapM* (**Table S4**).

These metabolically related polar lipids share a conserved glycosylated phosphatidylinositol (GPI) anchor and are integral components of the mycobacterial cell envelope. Disruptions in their synthesis enhance bacterial susceptibility to stress (e.g., treatment with ethambutol) while also affecting cell shape and division (28). Since RNA-Seq results suggested potential alterations in mycobacterial cell envelope lipid composition, we performed thin-layer chromatography (TLC) of lipid extracts, which revealed an enrichment of the PIMs fraction in the *ΔnapM* strain compared to the wild-type strain (**Fig. 5A**). Additionally, observed downregulation within the PG synthesis pathway, and to a lesser extent within the arabinogalactan (AG) synthesis pathway (**Table S4**), suggest a possible disruption of the cell wall skeleton structure, potentially impacting the integrity of the mycolic acid (MA) layer, which is anchored in PG-AG matrix. Therefore, we extended our analysis of cell wall lipids by TLC of mycolic acid methyl esters (**Fig. S4A**). However, no significant changes were observed, which is consistent with the RNA-Seq data showing no alterations in transcripts associated with MA biosynthesis.

**Figure 5.**
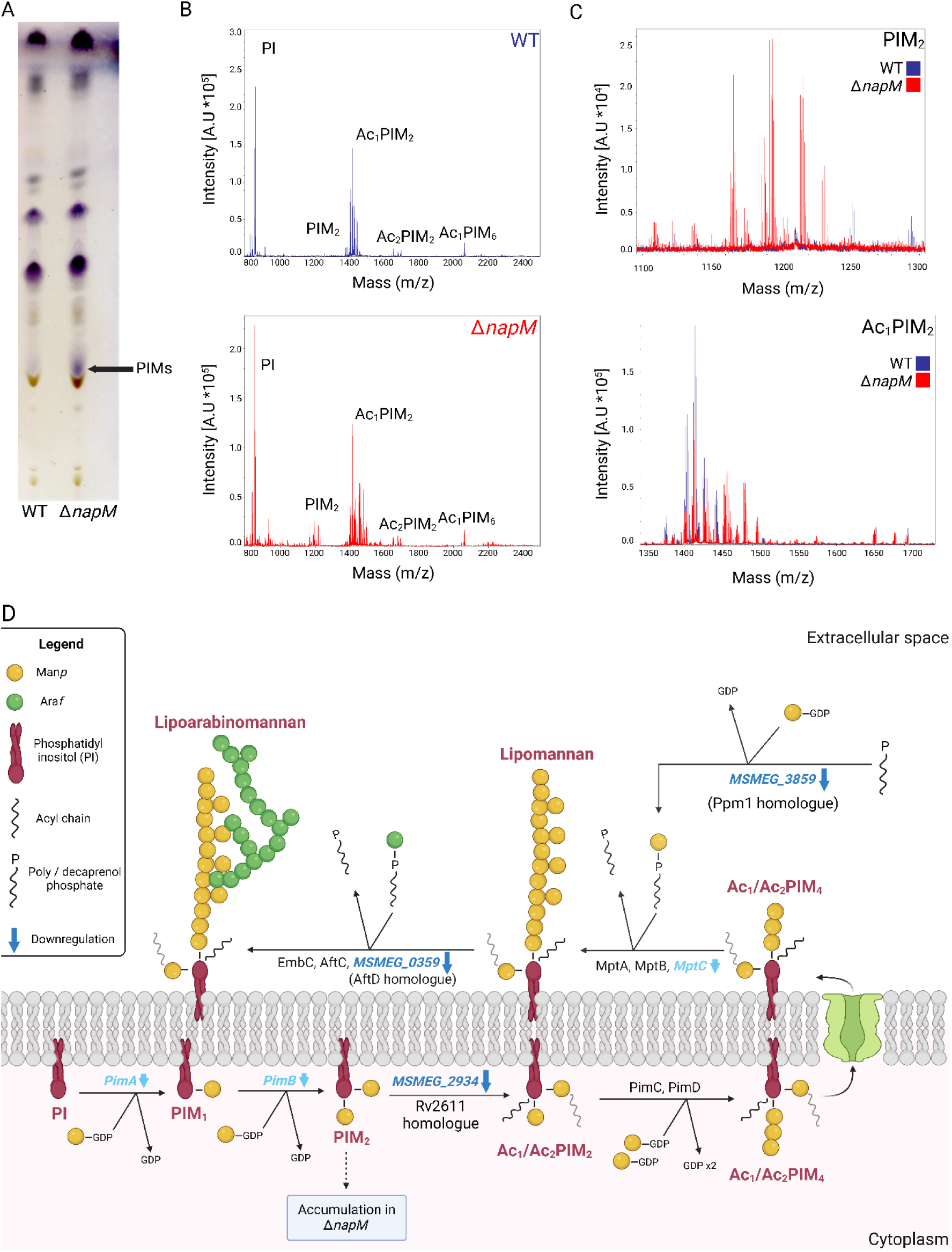
*napM* deletion results in altered PIMs composition. **A.** TLC analysis of the total lipid fractions of the wild-type (WT) and Δ*napM* strains. **B.** MALDI-TOF analysis of the wild-type (WT) and Δ*napM* strains. PIM_2_ is being accumulated in Δ*napM* (red) strain and the acylated PIM_2_ level is lowered in comparison to the wild-type strain (blue). **C.** Zoomed regions of MALDi-TOF spectra of PIM_2_ (phosphatidylinositol dimannoside) and Ac_1_PIM_2_ (acylated phosphatidylinositol dimannoside) obtained from the wild-type (blue) and Δ*napM* cells. **D.** Schematic biosynthesis pathway, with implemented transcriptomic changes of (blue – downregulated; red – upregulated) polar lipids (PI, LM and LAM) in *M. smegmatis* Δ*napM* strain. **E.** Transcriptomic changes in the inositol and phosphatidylinositol biosynthesis pathways in *M. smegmatis* Δ*napM* strain.

2D-TLC (**Fig. S4B, C**) of total lipids derived from the Δ*napM* strain confirmed the presence of an additional lipid species, later identified by MALDI-TOF analysis as non-acylated PIM_2_ (phosphatidylinositol dimannoside) (**Fig. 5B, C**). A similar PIM profile, characterized by the accumulation of PIM_2_ and a reduction in the acylated form of PIM_2_ (Ac_1_PIM_2_), was observed in *M. smegmatis* strain lacking *MSMEG_2934*, the gene that encodes the acyltransferase responsible for the addition of an acyl group to PIM_2_ (29). Indeed, in the Δ*napM* strain, *MSMEG_2934* is the most downregulated gene in the PIMs biosynthesis pathway, together with slight downregulation of the other genes encoding key acyltransferases, *pimA*, and *pimB* (**Fig. 5D**). RNA-Seq analysis reveals that in addition to significant changes in the expression of transcripts involved in phosphatidylinositol-containing lipids synthesis, other pathways associated with inositol utilization and inositol metabolism itself are also affected in the Δ*napM* strain (see **Table S4**). Notably, one of the most upregulated genes in the Δ*napM* strain is *ino1* gene (**Fig. 6**), which encodes inositol-1-phosphate synthase, a crucial enzyme for inositol synthesis (30). Interestingly, genes located in the ABC transporter gene cluster (*MSMEG_4658 - MSMEG_4656)*, previously described as essential for the uptake of exogenous inositol (31), are also highly upregulated in the Δ*napM* mutant. Inositol is not only essential for the synthesis of PIM, LM, and LAM (32) but also serves as a key precursor in the synthesis of mycothiol, a vital component of the cell’s antioxidant defense and detoxification pathways (30, 33) (**Fig. 6**). It is worth noting that, similar to the key genes involved in PIM, LM, and LAM synthesis, the major genes of mycothiol synthesis pathway are also downregulated when *napM* is deleted (**Table S4, Fig. 4**) (33).

**Figure 6.**
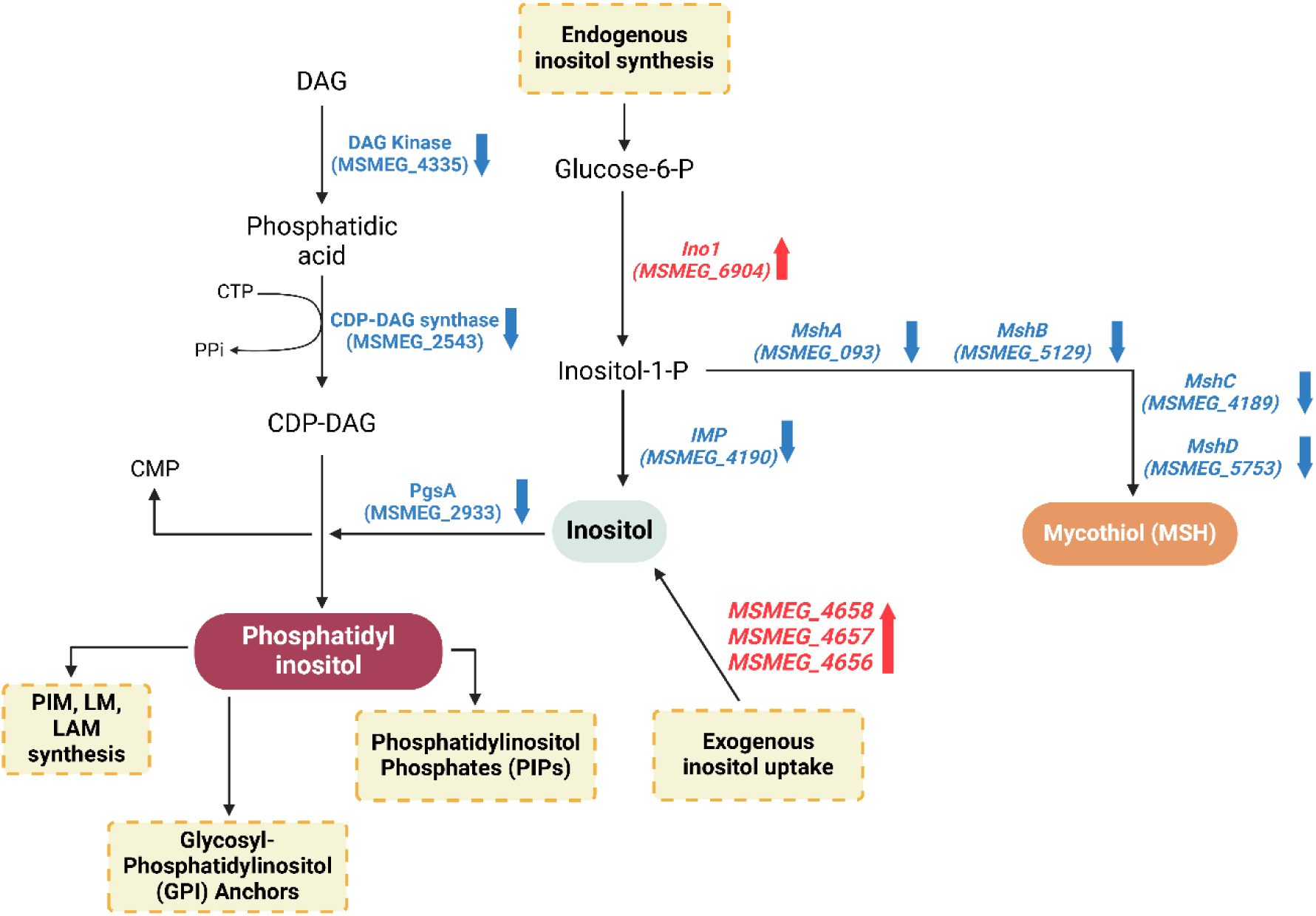
Transcriptomic changes in the inositol and phosphatidylinositol biosynthesis pathways in *M. smegmatis ΔnapM* strain. Blue indicates downregulation in the Δ*napM* strain, while red represents upregulation.

Our findings suggest that NapM may play a pivotal role in regulating cell envelope components synthesis, particularly by modulating PIMs biosynthesis, which are crucial for mycobacterial survival under stress conditions (34). By acting as an activator of genes involved in various polar lipid biosynthesis pathways, including PI, PIMs, LM, and LAM, NapM appears to shape the cell envelope structure, potentially enabling *M. smegmatis* cells to adapt to environmental challenges.

### NapM is a stress-induced protein

Our previous RNA-Seq experiments (14) suggest that the *napM* gene is expressed at a relatively low level in exponentially growing *M. smegmatis* cells, and data from the protein abundance database (PaxDb, https://pax-db.org/protein/246196/MSMEI_6719) indicate that NapM protein constitutes a minor fraction of total cellular proteins, with levels between 12.0 and 17.7 ppm, compared to 12,000–14,000 ppm for HupB. Intriguingly, the level of *napM* transcript in pathogenic *M. tuberculosis* increases upon exposure to various stresses, including environmental factors and antibiotics (22). However, no such information is available for NapM in saprophytic *M. smegmatis*, which occupies different, highly competitive environmental niches and possesses a larger genome than *M. tuberculosis*. We investigated the impact of *napM* deletion on *M. smegmatis* growth in response to stress. We found that *napM* deletion impacts *M. smegmatis* growth even under optimal conditions, as evidenced by a reduced growth rate in the lag phase without alteration in the exponential phase (**Fig. 7**). This appears to contrast with a previous report that concluded *napM* deletion in *M. smegmatis* had no impact on growth in optimal conditions using CFU measurements (21). However, the potential identification of dormant cells, which are not detected by growth curve analysis, may explain this discrepancy. We also examined growth of the *napM* deletion *(ΔnapM*) under stress conditions generated using the disinfectants (general environmental stressors) and antibiotics targeting different cellular processes (i.e., cell envelope biosynthesis, lipoarabinomannan biosynthesis, arabinogalactan biosynthesis, transcription, replication). The *ΔnapM* strain exhibited delayed adaptation and reduced growth compared to the WT strain when exposed to agents that directly target cell envelope integrity—such as vancomycin, ethambutol, benzalkonium chloride (BAC), nisin—and indirectly, like triclosan, underscoring the role of NapM in cell envelope resilience. In the presence of rifampicin, a transcription inhibitor, Δ*napM* cells exhibited significantly reduced growth. Unexpectedly, treatment with novobiocin, a gyrase B subunit inhibitor, resulted in enhanced growth of the mutant strain (**Fig. 7**) suggesting the NapM- mediated bypassing mechanism for maintaining proper chromosomal DNA topology under treatment with this antibiotic. For the NapM↑ strain, we observed a noticeable difference in growth only in the presence of vancomycin and nisin, with a significant reduction in growth compared to the WT strain (see **Fig. S5**).

**Figure 7.**
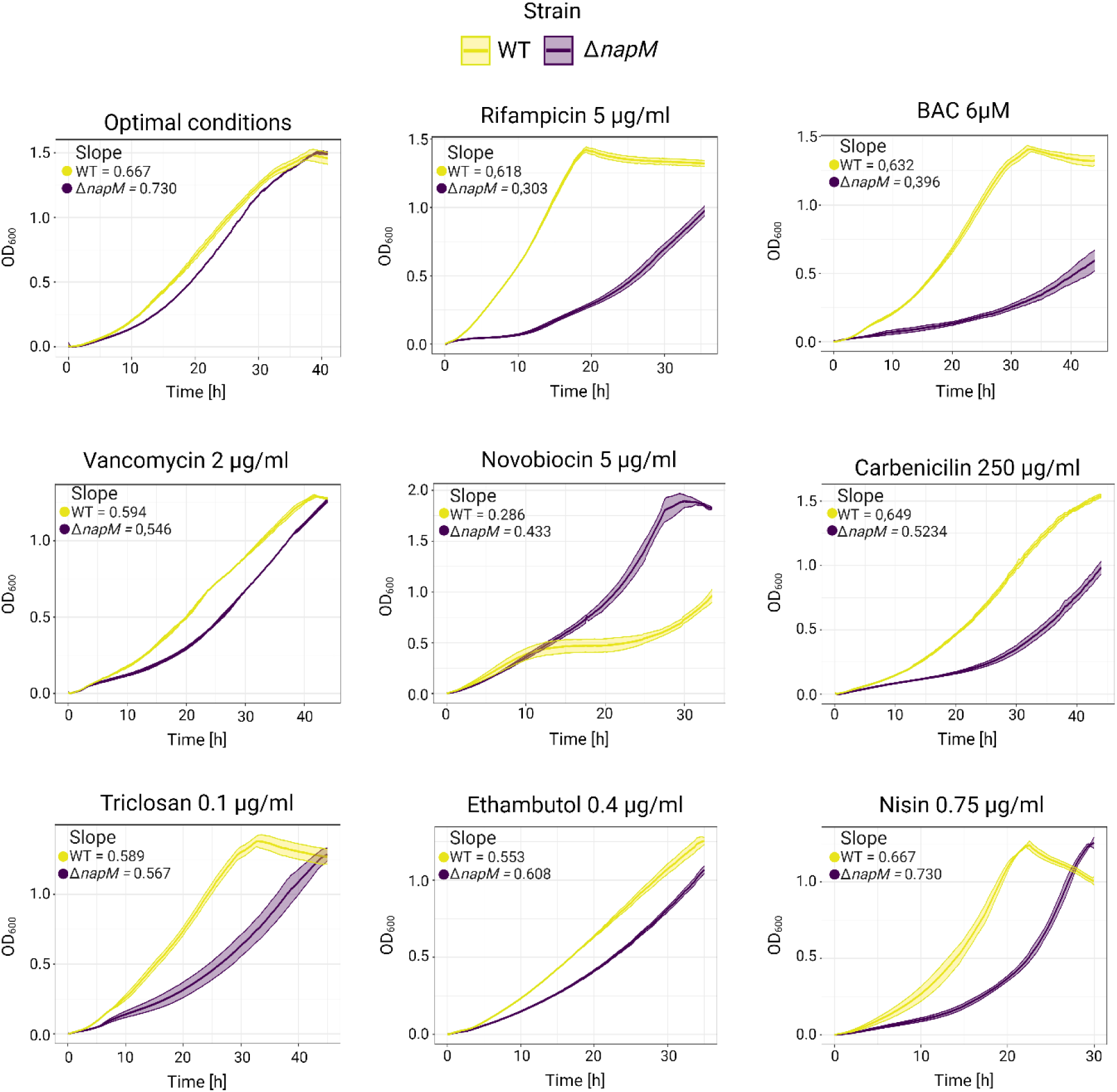
Loss of *napM* affects the growth of *M. smegmatis* under various stress conditions. Growth curves of the Δ*napM* strain compared to the *M. smegmatis* mc^2^ 155 wild-type strain (WT).

Additionally, we tagged NapM with FLAG and analyzed the fusion protein level during the growth of *M. smegmatis* and under various stress conditions. The NapM levels were elevated in exponentially growing *M. smegmatis* upon exposure to all tested stressors, with notable increases observed following treatment with ethambutol and disinfectant - BAC and triclosan (**Fig. S6**). These three stressors compromise the integrity of the cell envelope: BAC disrupts the cell membrane through direct physical damage; triclosan inhibits fatty acid biosynthesis, which indirectly affects membrane stability; and ethambutol inhibits arabinosyltransferases (EmbA-C), which disrupts the formation of the cell wall carbohydrates arabinogalactan and lipoarabinomannan.

In summary, NapM appears to play a pivotal role in mediating mycobacterial adaptation to stressors targeting the cell envelope (e.g., BAC, EMB, triclosan), transcriptional machinery (e.g., rifampicin), and DNA supercoiling (e.g., novobiocin).

### NapM localizes in the septum in response to stress

Given that NapM is a stress-induced DNA-binding protein, we investigated its subcellular localization under both optimal and stress conditions. Previous studies demonstrated that NapM colocalizes with chromosomal DNA in *E. coli* cells (21), a finding we confirmed in our experiments (**Fig. S7**). To analyze the localization of NapM during the *M. smegmatis* cell cycle, we constructed a strain with *napM* fused to the *mneongreen* gene at the native locus (NapM-mNeonGreen strain). Fluorescence microscopy of NapM-mNeonGreen cells under optimal conditions revealed a diffuse distribution of the fusion protein, with no distinct localization (**Fig. 8A, left panel**).

**Figure 8.**
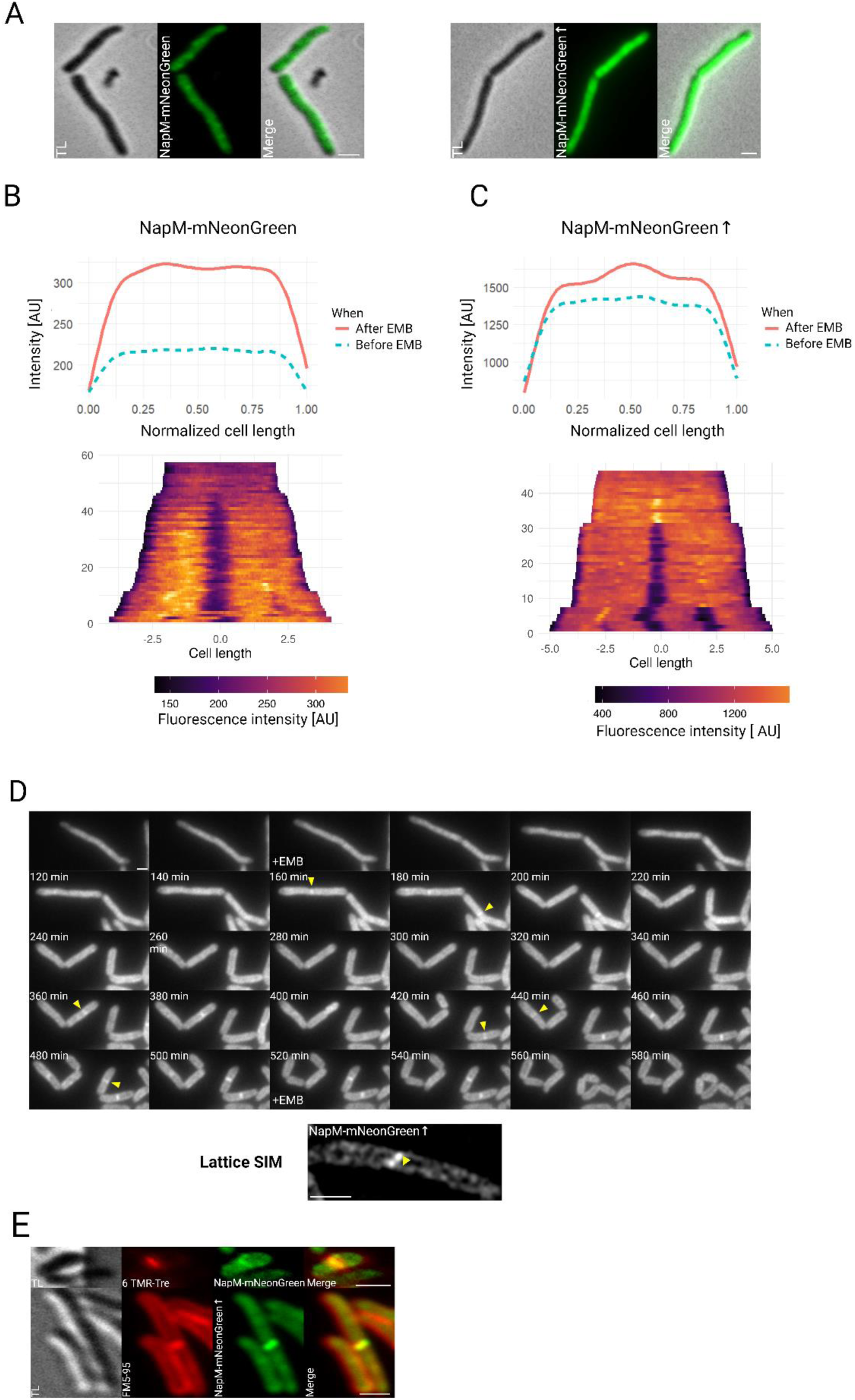
NapM localizes to the septum after treatment with ethambutol (EMB). **A.** NapM- mNeonGreen localization under optimal conditions. Scale bar, 1 µm. TL – transmitted light. **B.** Averaged fluorescence profile measured along the long axis of the cell (n = 100) and kymograph for a representative cell expressing NapM-mNeonGreen from the native locus and **C.** for cells overexpressing NapM-mNeonGreen (NapM–mNeonGreen↑). **D.** Time-lapse analysis of the *M. smegmatis* NapM-mNeonGreen↑ strain, showing localization of NapM- mNeonGreen at the septum area (yellow rectangles) after exposure to EMB. Scale bar, 1 µm. **E.** NapM-mNeonGreen and NapM-mNeonGreen*↑* cells stained with cytokinesis marker, FM5- 95, showing colocalization of NapM-mNeonGreen complexes with the formed septum after EMB treatment. The right panel presents high-resolution image of a representative NapM- mNeonGreen*↑* cell after EMB treatment, acquired utilizing Lattice SIM imaging. Scale bar, 1 µm. TL – transmitted light.

Interestingly, upon exposure to ethambutol (**Movie 1**), when the level of NapM is elevated (**Fig. S6**), the NapM-mNeonGreen fusion protein exhibited distinct localization at the newly formed septum. However, this phenomenon was observed in only a small fraction of dividing cells. Notably, mycobacteria exhibit cell-to-cell heterogeneity, particularly in response to environmental stress including antimicrobial compounds (35).

To further explore this specific septal localization, we constructed a merodiploid strain containing an additional *napM-mneongreen* fusion gene under the control of an inducible promoter (NapM-mNeonGreen↑ strain), while retaining *napM-mneongreen* at the native locus. This approach enabled us to examine whether elevated NapM levels (**Fig. S8, Fig. S9A**) would enhance or stabilize its septal localization. Although the NapM-mNeonGreen↑ strain displayed a stronger fluorescence signal compared to the native NapM-mNeonGreen strain, no distinct localization patterns were observed under optimal conditions (**Fig. 8A, right panel**). However, upon EMB treatment, fluorescence intensity markedly increased in both strains (**Fig. 8B**), and robust septal localization of NapM-mNeonGreen was evident in the majority of dividing cells in merodiploid strain (**Fig. 8C, Movie 2**). A similar NapM localization was also detected in response to triclosan exposure (**Fig. S10**). The NapM-mNeonGreen fusion protein was observed at the septum for approx. 40 – 50 minutes (40 ± 19.63 min for NapM- mNeonGreen and 50 ± 22.62 min for NapM-mNeonGreen↑, n = 87, **Fig S11A**) and disappeared immediately after division (i.e., V-snapping) (**Fig 8D upper panel**). High resolution microscopy using Lattice SIM further confirmed the septal localization of the NapM-mNeonGreen fusion protein following EMB treatment (**Fig. 8D, lower panel**). Additional experiments employing FM5-95 dye (a cytokinesis marker) and TMR- Trehalose dye (a mycolic acid marker) verified that NapM-mNeonGreen indeed colocalizes with the newly formed septum (**Fig. 8E**). Analysis of cells exhibiting septal localization of NapM revealed that NapM-mNeonGreen↑ cells were longer than their NapM-mNeonGreen counterparts (3.42 ± 0.77 µm vs. 2.86 ± 0.59 µm; t(198) = 5.77, p = 7.9 × 10⁻⁸, n = 100 per group; see **Fig. S11B**). Interestingly, these observations contrast with the previously reported phenotype of NapM↑ cells under optimal conditions, which were shorter than those of the Control↑ strain. Furthermore, we observed that NapM-mNeonGreen↑ cells with septal localization of NapM exhibited a greater degree of asymmetric division compared to NapM-mNeonGreen cells (**Fig. S11C**).

Taken together, these results show that the subcellular localization of NapM differs from that of previously characterized NAPs in *Mycobacterium*. Under optimal conditions, the NapM-mNeonGreen signal is diffused through the entire cell, with no discrete fluorescent foci. However, upon exposure to the cell envelope-targeting drugs, ethambutol, and triclosan, it localizes in the septum, suggesting that NapM might play a role in modulating cell division when encountering stressors that affect the integrity of the cell envelope.

## Discussion

Our data revealed that NapM, annotated as a PadR-family transcriptional regulator, is a highly conserved protein within the *Mycobacterium* genus (**Fig. S1**), and forms homodimers (**Fig. 1A**) - a feature supported by the recently resolved crystal structure of *M. tuberculosis* NapM (https://www.rcsb.org/structure/8JXK). While the level of NapM is significantly lower than that of other NAPs like HupB or mIHF, it increases markedly in response to cell envelope-targeting agents, such as benzalkonium chloride, ethambutol, and triclosan (**Fig. S6**). This observation suggests that NapM plays a crucial role in the adaptation of *M. smegmatis* to antimicrobial stress.

Consistent with this hypothesis, the absence of NapM had only a marginal effect on growth under optimal conditions, slightly prolonging the lag phase, but significantly impaired growth under stress conditions (**Fig. 7**). Interestingly, the *ΔnapM* strain displayed enhanced growth in the presence of novobiocin, gyrase B inhibitor. Although no changes were detected in the transcripts of *gyrB* (*MSMEG_005*) or *MSMEG_0457* (encoding the DNA topoisomerase IV subunit B), the upregulation of *MSMEG_1229*, annotated as a homolog of *gyrB* (which lacks in *M. tuberculosis*), may compensate the novobiocin action.

Unlike in *M. tuberculosis*, NapM in *M. smegmatis* does not appear to regulate chromosome replication at the initiation stage (**Fig. 3, Fig. S3**), as it presumably not interacts with the DnaA protein. Instead, RNA-Seq analysis (see **Table S4**) suggests that NapM indirectly inhibits DNA replication by modulating the transcription of the *dnaN* and *dnaQ* genes encoding the β and ε subunits of DNA polymerase III, respectively. Under nutrient-limiting conditions, the absence of NapM may disrupt the metabolic network, thereby impairing the stress response, as evidenced by the overall slowdown of the cell cycle (see **Fig. 3**). Hence, NapM is likely to orchestrate chromosome replication dynamics under stress by finely regulating the expression of genes essential for DNA synthesis and cellular metabolism.

Deletion of *napM* resulted in altered transcription of over 2, 000 genes (**Fig. 4A**), including those involved in cell envelope synthesis, particularly the biosynthesis of polar lipids such as PI, PIM, LM, and LAM (**Table S4** and **Fig. 5D**). Lipid profiling using TLC and MALDI-TOF revealed an enrichment of PIMs, especially PIM_2_, and a concomitant reduction in acylated PIM_2_ species (Ac_1_PIM_2_) in the *ΔnapM* strain compared to the wild-type strain (**Fig. 5A-C**). These findings are consistent with transcriptomic data showing downregulation of *MSMEG_2934*, which encodes an acyltransferase homologous to Rv2611 from *M. tuberculosis*. This enzyme catalyzes the acetylation of PIM_2_ to form Ac_1_PIM_2_/Ac_2_PIM_2_, precursors for more complex structures such as LM and LAM (36). Notably, deletion of *MSMEG_2934* in the wild-type strain produced a lipid profile (29) similar to that observed in the Δ*napM* strain (**Fig. 5, 6**).

Beyond polar lipids, genes involved in peptidoglycan (PG) biosynthesis and cell division were also dysregulated in the *ΔnapM* strain (**Table S4**), highlighting the role of NapM in coordinating cell envelope biosynthesis and division under stress. These disruptions likely explain the elongated cell morphology observed under optimal conditions (**Fig. 2A**) and the growth defects exhibited by Δ*napM* cells when exposed to cell envelope-targeting stressors (**Fig. 7**).

Contrary to its initial classification as a NAP, NapM did not exhibit distinct nucleoid localization under optimal growth conditions in *M. smegmatis* cells (**Fig. 8A**) and did not affect the nucleoid-to-cell length ratio in Δ*napM* cells (**Fig. S2B**). However, the nucleoid was less compacted (i.e., elongated) in *ΔnapM* cells (**Fig. 2D**). This phenotype may result from reduced molecular crowding and weaker depletion forces in elongated cells, or from NapM potential role in modulating chromosome architecture. Remarkably, under exposure to ethambutol (**Fig. 8C-E**) or triclosan (**Fig. S10**), NapM displayed unique septal localization in dividing cells (**Movie 1**, **Movie 2**) - a phenomenon not reported for any other NAP studied to date.

Given its stress-induced septal localization (**Fig. 8**) and RNA-Seq data showing that NapM regulates genes encoding cell envelope components (**Figs 4** and **5**, **Table S4**), we propose that NapM influences cell division process through a dual mechanism. In addition to activating cell envelope-related genes, NapM may interact with divisiome components, thereby facilitating septum formation and protecting chromosomes from guillotining. Supporting this hypothesis, NapM upregulates key division-associated genes, such as *wag31* gene (encoding a DivIVA homolog) (37, 38) *crgA* (encoding CrgA, which interacts with DivIVA) (39), and *ssd* (*MSMEG_6171*), all of which are essential for polar growth and septum formation (40).

Although NapM was initially perceived as a NAP, our findings characterize it as a pleiotropic regulator that modulates cell envelope composition, cell division, and chromosome replication dynamics. By modulating the transcription of critical genes, NapM enables *M. smegmatis* adapt to environmental stress and preserve cellular integrity. However, its lack of chromosomal localization, its unique stress-induced septal localization, and its involvement in polar lipid biosynthesis challenge its designation as a NAP (**Fig. 9**). These findings raise an intriguing question: if NapM is not a true NAP, how should this multifaceted protein be classified?

**Figure 9.**
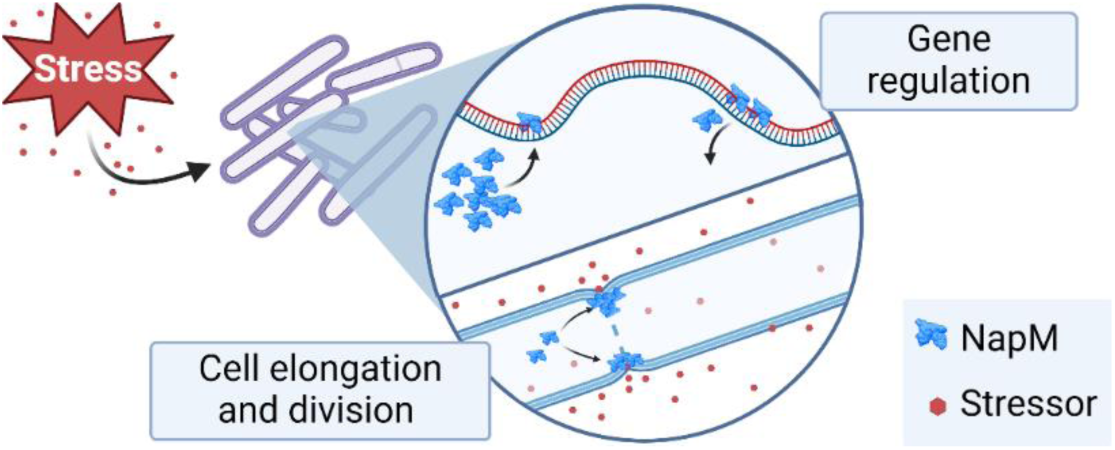
A graphical visualization depicting the role of NapM in *M. smegmatis* as both a stress-induced regulator and potential component of the divisome.

## Materials and methods

### RNA isolation

Total RNA was isolated as previously described (41, 42). Cells were cultured in 50 ml to OD_600_ 8 - 1.2, pelleted, resuspended in 300 μl DEPC H_2_O, transferred to a clean Eppendorf tube, and mixed with 900 μl TRIzol LS (Invitrogen) at a 1:3 ratio. After being briefly vortexed, the mixture was transferred to a new tube with zirconia beads compatible with the MP FastPrep24 homogenizer (VWR). The sample was cooled on ice for 5 minutes and then homogenized (2×45 sec; 6 m/s) with intermittent cooling on ice. The lysate was cleared by centrifugation (12,000 x g, 5-7 minutes, 4°C), and the supernatant was transferred to a new Eppendorf tube. To obtain the water-soluble fraction, the supernatant was incubated for 5 minutes at room temperature in a laminar flow hood. Chloroform (250 μl) was added and the mixture was vortexed, placed on ice, mixed vigorously for ∼ 2 minutes, incubated on ice for 10 minutes, and centrifuged (12,000 x g, 4°C, 15 min). The aqueous phase (upper layer) was transferred to a new Eppendorf tube and supplemented with isopropanol (1V), sodium acetate (1:10, 5M, pH 5.2), and Glyco Blue co-precipitant (Thermo Fisher Scientific), the mixture was vortexed, and RNA was precipitated for two days at −80°C. The tube was centrifuged (20,000 x g, 4°C, 30 min), and the precipitated RNA was resuspended with 500 μl of 70% ethanol (prepared with DEPC-treated water) and pelleted by centrifugation (20,000 x g, 4°C, 30 min). The obtained RNA pellet was air-dried in an open Eppendorf tube, suspended in 50-55 μl DEPC-treated water, and either stored at −80°C or immediately subjected to DNase digestion. DNase digestion was performed using 10 μg of RNA in a total volume of 50 μl at 37°C for 15 minutes with DNase I (A&A Biot). The RNA was then purified using magnetic beads (MagBind), eluted with 30 µl DEPC-treated water supplemented with RiboLock RNase inhibitor (Thermo Fisher Scientific), and stored at −80°C for further procedures.

### RNA-Seq

Total RNA of each analyzed strain was subjected to rRNA depletion utilizing the Pan-Actinobacteria riboPOOL (siTOOLs BIOTECH) and its corresponding protocol. Subsequently, the KAPA RNA HyperPrep kit (KK8544, ROCHE) was employed following the manufacturer’s instructions. Illumina system-compatible adapters, incorporating sample-specific 8-nucleotide-long barcoding sequences, were ligated, and cDNA libraries were PCR amplified. The resulting sequencing libraries were assessed on an Agilent 2100 Bioanalyzer with a DNA 1000 chip and quantified through real-time PCR using the NEBNext Library Quant kit for Illumina (New England Biolabs). Raw sequencing data were generated on a NextSeq500 platform (Illumina) using paired-end 75-cycle sequencing run-compatible reagents (150 cycles, NextSeq 500/550 Mid Output v2 sequencing kit; Illumina). Biological triplicates were prepared and sequenced for each growth condition.

For analysis of RNA-seq data, the initial processing involved demultiplexing and removal of library adapters (implemented with Cutadapt v.1.3) (43). A filtering step was used to exclude short reads (<20 bp) and those with low quality (<30%) using Sickle script v.1.33 (44)The subsequent alignment of high-quality reads to the *M. smegmatis* mc^2^ 155 genome (retrieved from NCBI, accession number NC_008596) was performed using the Bowtie2 short-read aligner (45). The resulting BAM files, containing read-mapping coordinates, were indexed and sorted with SAMtools v.1.7 (46) for data visualization (Integrative Genomics Viewer (44) and mapping of read counts to gene features (HTSeq.count script (47)). The read counts from individual samples were consolidated into a count matrix and submitted to the online differential expression RNA-seq analysis platform, Degust (http://doi.org/10.5281/zenodo.3501067). The analysis utilized default parameters and the voom/limma method (48) was selected for final data evaluation. Visual representation of the differential expression data was achieved through the creation of a volcano plot and heatmaps, using the Python Seaborn software suite (https://pypi.org/project/seaborn). Raw sequence reads and processed data (count matrix) were submitted to data repository GEO (NCBI) and is available under this accession (GSE275852).

### Fluorescence microscopy

To visualize NapM in *E. coli* cells, the PCR-amplified product, generated using primers pACYC_napM_Fw and pACYC_mNG_NheI_Rv (**Table S1**), was cloned into the NcoI and HindIII sites of the pACYCDuet™-1 vector (Sigma). Transformants were selected on LB agar supplemented with chloramphenicol. The resulting plasmid was verified by PCR and sequencing, and transformed into *E. coli* BL21(DE3) cells. Positive clones were used in fluorescence microscopy experiments. An overnight culture of the transformants was grown in LB liquid medium supplemented with chloramphenicol and a small portion of the culture was used to inoculate fresh medium. Once the culture reached an OD_600_ of 0.4–0.6, isopropyl β-D-1-thiogalactopyranoside (IPTG) was added to a final concentration of 1 mM, and the culture was incubated for 1 hour. Subsequently, 1 ml of the culture was stained with the chromosomal dye DAPI (Molecular Probes) at a final concentration of 2 µg/ml for 20 minutes. Cells were centrifuged at 5000 × g for 5 minutes, washed with phosphate-buffered saline (PBS), and resuspended in PBS. The suspension was smeared onto microscopic slides. As a negative control, *E. coli* BL21(DE3) cells transformed with an empty pACYCDuet™-1 vector were used.

To visualize NapM fluorescence fusion proteins, snapshot analysis was performed on exponential-phase *M. smegmatis* cells (OD600 = 0.8–1.2). Overnight cultures of *M. smegmatis* were grown in 7H9 medium supplemented with ADC and 0.05% Tween 80. The culture was centrifuged at 5000 × g for 5 minutes at room temperature, and the pellet was washed once with PBS and resuspended in PBS. For cell membrane visualization, log-phase cells (OD600 = 0.8–1.2) were stained with FM5-95 dye (0.5 µg/ml; Thermo Fisher Scientific). For cell wall visualization, 6-TMR Tre (100 µM; Torcis Bio-Techne) was used, and for chromosomal staining, Hoechst 33342 (1 µg/ml; VWR) was applied. In each case, 1 ml of cells was incubated with the dye for 30 minutes at 37°C with continuous agitation at 180 rpm, followed by centrifugation at 5000 × g for 5 minutes at room temperature. The pellet was then washed with PBS and the cells resuspended in PBS. All samples were mounted on Teflon-coated glass slides with 1.2% agarose and examined using a Leica DM6 epifluorescence microscope equipped with a 100×/1.4 oil objective and appropriate filters (DAPI, GFP, mCherry). Images were analyzed using Fiji software (49) and R software (R Foundation for Statistical Computing, Austria; http://www.r-project.org) with the ggplot2 package.

### Time-lapse fluorescence microscopy (TLFM)

Overnight liquid cultures of *Mycobacterium smegmatis* (OD_600_ ∼ 0.5) were observed using an ONIX microfluidics system. Briefly, 70 µl of bacterial culture was introduced into the cell inlet well of the ONIX B04A plate (Merck). For each experiment, the first two wells were flushed with PBS, and 150–300 µl of 7H9 medium supplemented with ADC and 0.05% Tween80 was loaded into each well. Once the cells were loaded into the observation chamber, they were washed for 45 minutes with the medium under 3 psi pressure. Subsequently, the cells were cultivated for 24 hours under 1.5 psi. Where required, cells were exposed to antibiotics for 6 hours under 1.5 psi, after which the antibiotic was removed by washing with the previously used medium. To induce nutrient-restricted conditions, the flow of the medium was halted following the washing step. Images were captured automatically at 10-minute intervals using a Delta Vision Elite inverted microscope equipped with a UPlanFL N 100×/1.3 Oil Ph3 objective and an environmental chamber set to 37°C. Time-lapse images were analyzed using Fiji software (49) and R (R Foundation for Statistical Computing, Austria; http://www.r-project.org), with data visualization performed using the ggplot2 package.

### Lattice SIM

Exponential-phase cells (OD_600_ = 0.8–1.2) exposed to ethambutol were imaged using an Elyra 7 (Zeiss) inverted microscope equipped with an sCMOS 4.2 CL HS camera and an alpha Plan-Apochromat 100×/1.46 Oil DIC M27 objective, along with an Optovar 1.6× magnification changer. Fluorescence was excited using a 488 nm laser (100 mW), and signals were captured through a multiple-beam splitter (405/488/561/641 nm) and corresponding laser-block filters (405/488/561/641 nm). Samples were prepared on agar pads (1% agarose in Milli-Q water, poured into 1.0 × 1.0 cm GeneFrames; Thermo Fisher Scientific). Cells were illuminated with a 488 nm laser at 5% intensity and a 30 ms exposure time in Lattice-SIM mode, consisting of 15 phases. Image reconstruction was performed using ZEN 3.0 SR software (Zeiss) with standard parameters. Images were analyzed using Fiji software (49) and R (R Foundation for Statistical Computing, Austria; http://www.r-project.org), with the ggplot2 package utilized for data visualization.

### Bacterial two-hybrid system

To verify the dimerization of Mycobacterium smegmatis NapM, we employed the Bacterial Adenylate Cyclase Two-Hybrid System (BACTH System Kit, EUROMEDEX). Briefly, derivatives of pUT18, pUT18C, pKT25, and pKNT25 were generated through PCR amplification using primers BTH_xbaI_napM_Fw and BTH_kpnI_napM_Rv (**Table S1**). The resulting products were initially cloned into the pGEM-T Easy vector (Promega) and then subcloned into backbone vectors through restriction cloning. The experimental procedures were conducted according to the manufacturer’s guidelines provided in the BACTH System Kit. Transformants were plated on LB agar supplemented with IPTG, kanamycin, ampicillin, and X-gal, and incubated for 2 days at 30°C. Plate images were captured using a Chemi Doc MP (Bio-Rad).

### Total lipid extraction and TLC analysis

Total lipids were extracted from dry (50 mg) and wet (400 mg) cell masses of Mycobacterium smegmatis wild-type and Δ*napM* strains. Each collected cell pellet was dissolved in water and subjected to chloroform-methanol (1:2, v/v) extraction. The combined extracts were partitioned in a chloroform-methanol-water mixture (1:2:0.8, v/v/v), and the organic phase was collected and dried at 40°C under a stream of nitrogen.

The obtained total lipids were dissolved in chloroform to a final concentration of 50 mg/ml. Equal amounts of lipid extracts were applied to silica gel 60-precoated HPTLC or TLC plates (Merck; layer thickness, 0.2 mm). Lipids were analyzed using TLC with a chloroform-methanol-water (60:30:6, v/v/v) solvent system. For visualization, TLC plates were sprayed with orcinol, vanillin, and iodine vapor, then heated at 120°C.

### MALDI-TOF

MALDI-TOF mass spectrometry was performed using an ultrafleXtreme spectrometer (Bruker). Spectra were measured in reflectron negative ion mode with the following parameters: ion source 1, 20.0 kV; ion source 2, 17.95 kV; lens, 6.5 kV; reflector, 21.10 kV; and reflector 2, 10.72 kV. The mass range analyzed was 700–3500 Da, using flexControl 3.4 software (Bruker). Approximately 2000 laser shots were totaled from one spot. The flexAnalysis 3.4 software (Bruker) was used for data analysis. Norharman (β-carboline) was used as the matrix.

### Statistical Analysis and data visualization

Each experiment was conducted with three biological replicates for each strain. Levene’s test was performed in RStudio (R Foundation for Statistical Computing, Austria; http://www.r-project.org) using the *car* package to assess variance equality. Depending on the results, either Student’s t-test (for equal variances) or Welch’s t-test (for unequal variances) was applied. Statistical analyses and visualizations were carried out in RStudio using the *ggplot2* and *stats* packages (R Foundation for Statistical Computing, Austria; http://www.r-project.org). Additional data visualizations were created using BioRender (https://www.biorender.com/). Significance levels were defined as follows: ns (not significant), ** – p < 0.01 *** – p < 0.001; **** – p < 0.0001.

## Acknowledgments

The authors are grateful to Agnieszka Strzałka for the assistance in RNA-Seq data analysis and Tomasz Łebkowski for critically reviewing the manuscript.

## Author contributions

JZ-C, JH, KM contributed to the conception and design of the study. KM, YL, PP optimize and performed RNA-Seq experiments and KM, PP, and JP analyzed RNA-Seq data. KM, JH, and JZ-C wrote the original draft. JP, MP, PP reviewed and edited the manuscript. KM performed analysis, and visualized data. KM, MP, YL, PP and JH performed experiments. JZ-C delivered resources and raised funds. All authors contributed to the article and approved the submitted version.

## Materials & Correspondence

Materials and correspondence should be addressed to Jolanta Zakrzewska-Czerwińska or Joanna Hołówka.

## Data availability statement

The original contributions presented in the study are included in the article/Supplementary Material. RNA-Seq data were deposited in the Gene Expression Omnibus (GEO), a public repository for functional genomics data, and are available under accession number #GSE275852. Further inquiries can be directed to the corresponding authors.

