## Supplementary data for "The mycobacterial nucleoid-associated protein NapM exhibits stress-induced septal localization and modulates cell envelope gene expression"

**Text S1. Supplementary Experimental Procedures**

### Plasmid Propagation and Bacteria Cultivation

Plasmids utilized for *Mycobacterium smegmatis* mc$^2$ 155 transformation were propagated in *Escherichia coli* DH5α. *E. coli* was cultured in Luria-Bertani (LB) broth at 37°C with agitation at 180 rpm and on LB agar plates (Difco) at 37°C (1). Media were supplemented with antibiotics (100 µg/ml ampicillin, 50 µg/ml kanamycin) or other additives if needed, including 0.004% X-Gal [5-bromo-4-chloro-3-indolyl-α-D-galactopyranoside]. *M. smegmatis* liquid cultures were grown in Middlebrook 7H9 supplemented with 0.05% Tween 80 and albumin-dextrose-catalase (ADC; BD) medium, or in Difco Nutrient Broth (NB; BD) medium at 37°C with agitation at 180 rpm. Solid medium cultivation involved the addition of 2% agar to NB or Middlebrook 7H10 medium with oleic acid-albumin-dextrose-catalse (OADC BD). Cultures were incubated at 37°C until visible colony formation occurred over 2-5 days. Antibiotics (100 µg/ml ampicillin, 50 µg/ml kanamycin, 2.5–7.5 µg/ml rifampicin, 0.4 µg/ml ethambutol, 5 µg/ml novobiocin, 6 µM BAC, 0.75 µg nisin, triclosan 0.1 µg/ml, vancomycin 2 µg/ml, carbenicillin 250 µg/ml) and other additives (0.004% X-Gal, 2% sucrose,1 mM Isopropyl β-D-1-thiogalactopyranoside (IPTG), 1% acetamide were added if required).

### Construction of *M. smegmatis mc$^2$* 155 mutant strains

The allelic replacement of *napM* gene *(MSMEG_6903)* with fusion genes *napM-mneongreen or napM-flag* was performed following the protocol outlined by Parish and Roberts (2). Briefly, the chromosome of *M. smegmatis mc$^2$ 155* was used as template to amplify downstream and upstream regions of *napM* gene using two primer sets napM1_FW x napM1_RV and napM2_FW x napM2_RV. In the case of *mneongreen* fusion gene, it was amplified by PCR utilizing primer sets - mNG_FW x mNG_RV. Amplified products were cloned to p2NIL Ø plasmid and transformed colonies were spread on LB medium supplemented with kanamycin. In the case of *napM-flagx3* fusion, *flagx3* complementary oligonucleotides (Genomed) - HindIII_Flagx3_FW x NheI_Flagx3_RV with HindIII and NheI overhangs were annealed and cloned to p2NIL *napM-mneongreen* digested with HindIII and NheI restriction enzymes.

Analogous cloning strategy was used for construction of *M. smegmatis mc$^2$ 155 strain* with deletion of *napM* gene (*MSMEG_6903*). Flanking regions of *MSMEG_6903* were amplified using primers sets - napM1_FW x napM1del_RV and napM2del_FW x napM2_RV. Obtained products were cloned to p2NIL Ø plasmid and transformants were plated on LB supplemented with kanamycin.

In the case of $pMV_{pAMI}$ Ø integrative plasmids carrying fusions of *napM* gene, the following primers were used- $pMV_{pAMI}$_napM_FW x $pMV_{pAMI}$_napM_RV and $pMV_{pAMI}$_mNG_FW x $pMV_{pAMI}$_mNG_RV for *napM-mneongreen*. Amplified products were cloned using SLIC (3) to $pMV_{pAMI}$ Ø plasmid. Obtained derivatives of $pMV_{pAMI}$ Ø plasmid were used to transform electrocompetent *M. smegmatis* cells. Transformants were spread on NB medium supplemented with kanamycin and colonies were verified by PCR.

All p2NIL and $pMV_{pAMI}$ derivatives were verified by sequencing (Microsynth). In the final step, *goal* cassette was cloned into PacI site of every p2NIL derivative. Electrocompetent *M. smegmatis* cells were electroporated with 0,5 µg – 8000 µg of NaOH/EDTA-treated plasmid DNA and unmarked mutants were selected according to the procedure described previously by Parish and Roberts (2). In short, after transformation, cells were plated on NB supplemented with kanamycin and X-gal. Blue single crossing-over (SCO) mutants were further spread for biomass (on NB plates supplemented with 2% sucrose and X-gal) which was then induced to second crossing-over. Chosen double crossing-over (DCO) mutants were verified by PCR and sequencing. If applicable, Western blotting was used to prove the production of fusion proteins.

To compare growth of the constructed *M. smegmatis mc$^2$ 155* strains, cells were grown at 37˚C in a final volume of 300 µl 7H9 (supplemented with ADC and 0.05% Tween 80, and 1% acetamide if applicable). Optical density measurements were taken at 20 min intervals for 30 – 60 h using a Bioscreen C instrument. To determine the differences in growth rates of the analyzed strains, $OD_{600}$ values within the linear range of the growth curves were log10-transformed, and the slopes of the resulting curves were used to compare the growth rates.

**Western blotting**

Western blotting analysis was performed to assess the levels of fusion proteins in NapM-FLAGx3, NapM-mNeonGreen, and NapM-mNeonGreen↑ strains under various conditions. The cultures with $OD_{600}$ = 0,8 were centrifuged, and the pellet was sonicated in PBS to prepare cell lysates. For SDS-PAGE 20 - 100 µg of the total protein of each lysate were used. Western blotting was performed by transferring proteins to a nitrocellulose membrane. The membrane was then blocked with 5% non-fat dry milk in TBST (TBS + 0.1% Tween-20) overnight at 4°C to prevent non-specific binding. The primary antibodies used were an anti-FLAG antibody (Sigma, dilution 1:1000) for NapM-FLAGx3 and an anti-mNeonGreen antibody (ChromoTek, dilution 1:1000) for NapM-mNeonGreen and NapM-mNeonGreen↑ strains. The membranes were incubated with the primary antibody for 1 h at room temperature. After washing with TBST, membranes were incubated with goat anti-mouse IgG secondary antibody conjugated with HRP (Invitrogen, dilution 1:5000) for 1 h at room temperature. Protein bands were visualized using Pierce™ SuperSignal™ West Pico PLUS Chemiluminescent Substrate (Thermo

Scientific) and detected with a chemiluminescence imaging system (ChemiDoc MP, Bio-Rad). Loading control was performed using Ponceau S Staining solution accordingly to the procedure provided by the manufacturer (ThermoFisher Scientific).

**Two-dimensional thin-layer chromatography (2D-TLC)**

2D TLC was performed with 300 µg of total lipids for each strain on HPTLC plates, which were developed in two directions: the first in the solvent system containing chloroform, methanol and water (65:25:4, v/v/v) and the second containing chloroform, acetic acid, methanol and water (80:15:12:4, v/v/v/v). The HPTLC plates were sprayed with vanillin reagent, which enabled the visualization of lipids (4).

**TLC of mycolic acid methyl esters (MAMES)**

Mycolic acids were obtained from dry cell mass by an alkaline method using 15% tetrabutylammonium hydroxide and then analyzed by TLC (4). The mycolic acid methyl esters were dissolved in chloroform in the concentration of 50 mg/ml and applied on silica gel 60 TLC plates (Merck). Lipids were analyzed by TLC in a solvent system: hexane-diethyl ether (85:15, v/v) and were visualized by 10% molybdophosphoric acid in an ethanol solution, followed by heating at 120°C (4).

**Table S1. Oligonucleotides used in this study –** see Excel file “Table Supplementary”

**Table S2. Plasmids used in this study –** see Excel file “Table Supplementary”

**Table S3. Strains used in this study**

| Name | Relevant genotype | Source |
|---|---|---|
| WT | *M. smegmatis* $mc^2$ *155* | Lab collection |
| Control↑ | *M. smegmatis* $mc^2$ *155 attBL5::*pMV306$_{pAMI}$ Ø | Lab collection |
| DnaN-mCherry | *M. smegmatis* $mc^2$ *155 dnaN-mcherry* | Lab collection |
| NapM-mNeonGreen | *M. smegmatis* $mc^2$ *155 napM-mneongreen* | This study |
| Δ*napM* | *M. smegmatis mc2 155* Δ*napM* | This study |
| NapM-mNeonGreen↑ | *M. smegmatis* $mc^2$ *155 napM-mneongreen, attBL5::*pMV306$_{pAMI}$ *napM-mneongreen* | This study |
| NapM↑ | *M. smegmatis* $mc^2$ *155 attBL5::*pMV306$_{pAMI}$ *napM* | This study |
| Δ*napM*/DnaN-mCherry | *M. smegmatis* $mc^2$ *155* Δ*napM, dnaN-mcherry* | This study |
| NapM-FLAGx3 | *M. smegmatis* $mc^2$ *155 napM-flagx3* | This study |
| NapM-mNeonGreen$_{Ec}$ | *E. coli BL21 (DE3)* pACYC *napM-mneongreen* | This study |
| pACYC Ø | *E. coli BL21 (DE3)* pACYC Ø | This study |

**Table S4. Comparative analysis of gene expression changes for polar lipid metabolism, mycothiol, and peptidoglycan biosynthesis in *M. smegmatis* Δ*napM* strain –** see Excel file “Table Supplementary”.

### Supplementary Figures

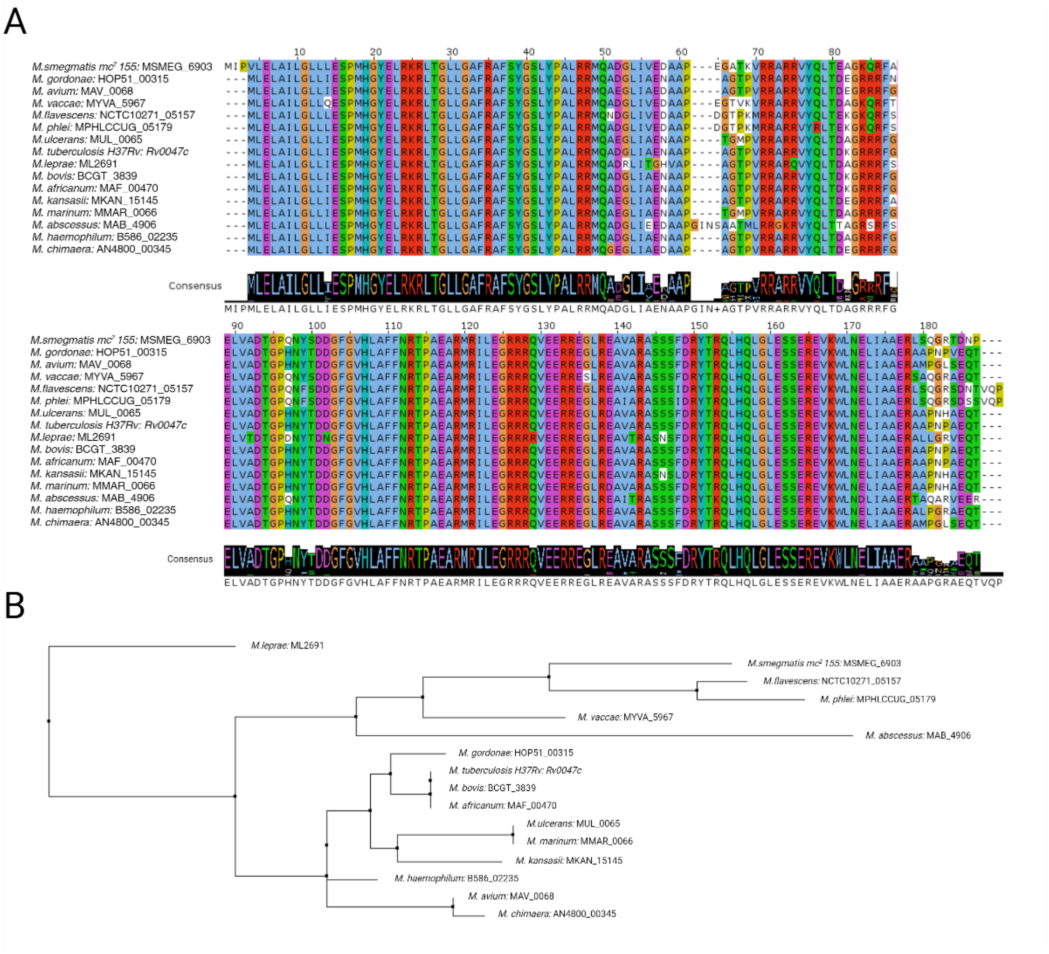


**Figure S1. NapM sequences across *Mycobacterium* species predicted with JalView software. A.** Multiple sequence alignment (Muscle) of mycobacterial NapM proteins. Upper panel – N-terminal domain; lower panel – C- terminal domain. **B.** Phylogenetic tree of NapM proteins in saprophytic and pathogenic mycobacterial species.

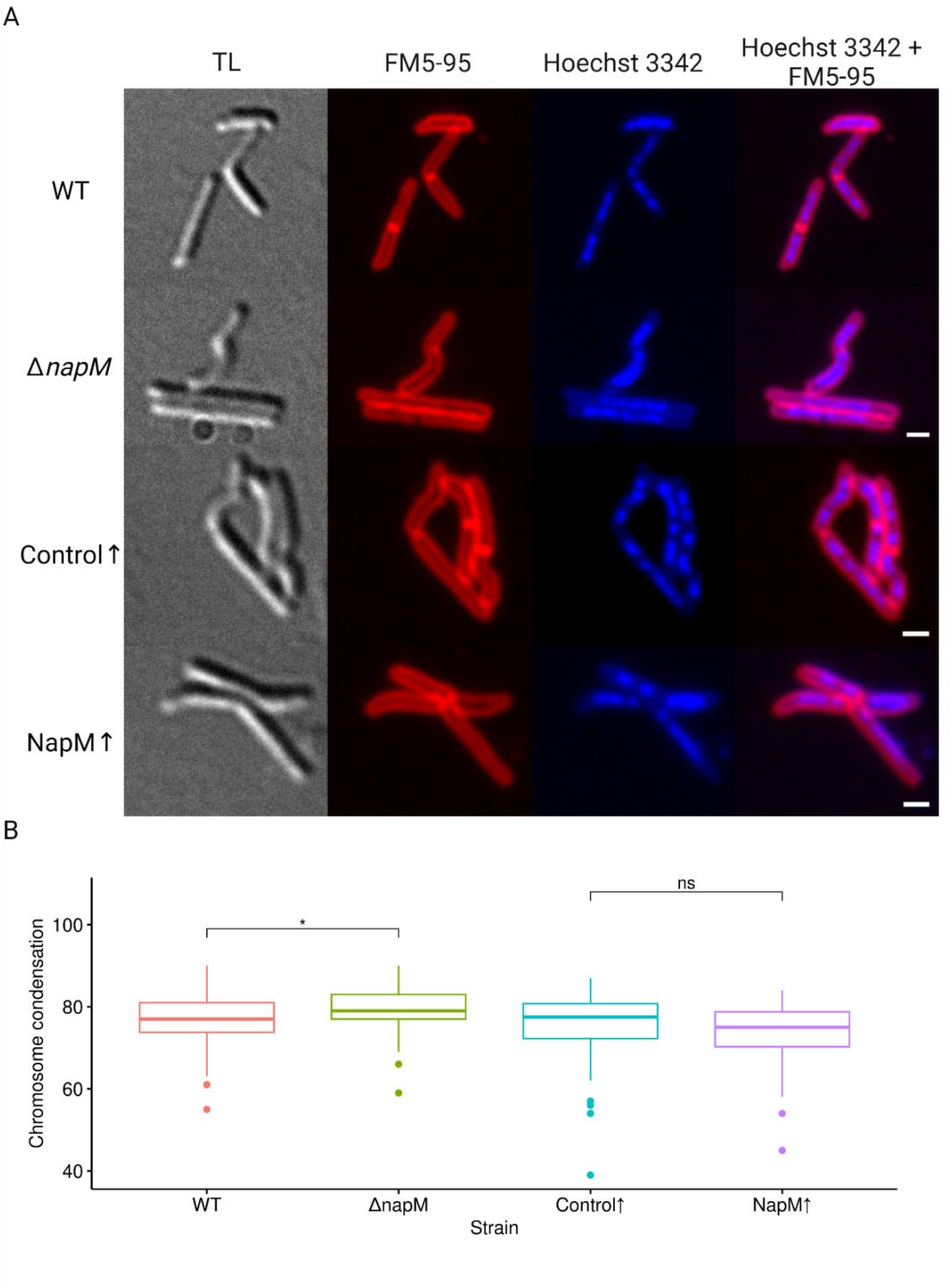


**Figure S2. Phenotypic analysis of the wild-type (WT) and Δ*napM* strains. A.** Membrane (FM5-95) and nucleoid (Hoechst 3342) staining of the analyzed *M. smegmatis* strains. TL – transmitted light. Scale bar, 1 µm. **B.** Chromosome condensation calculated as the ratio of chromosome length to the cell length. In the Δ*napM* cells, chromosome is slightly less condensed in comparison to the wild-type cells (76.87 ± 6.79 and 79.37 ± 5.50, respectively; $t(98) = -2.02$, $p = 4.2*10^{-2}$; $n = 50$), while chromosome condensation in NapM↑ cells is similar to the Control↑ cells (73.58 ± 7.28 and 74.68 ± 9.62, respectively; $t(98) = -0.64$, $p = 5.1 \times 10^{-1}$; $n = 50$).

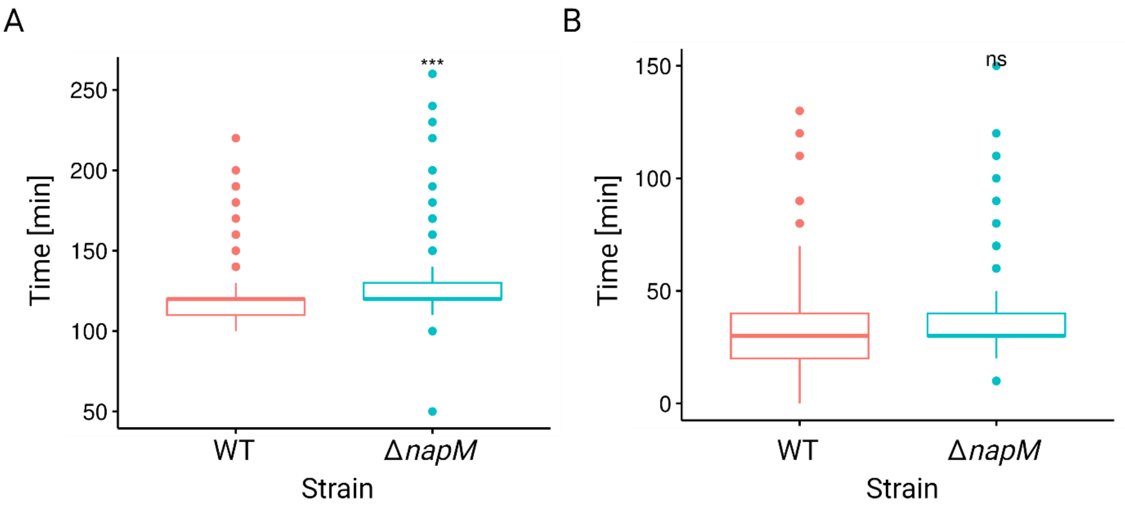


**Figure S3. Analysis of replication dynamics in Δ*napM* and the wild-type (WT) strains under optimal conditions. A.** Boxplot presenting duration of replication (C phase) in Δ*napM* and the wild-type strain (WT). Deletion of *napM* gene results in slight elongation of the C phase (replication) in *M. smegmatis* (125.58 ± 21.30 min and 120.20 ± 16.84 min, respectively; $t(608) = 3.46$, $p = 5.4*10^{-4}$, $n = 305$). **B.** Boxplot presenting the time between termination of replication and initiation of replication in daughter cells (BD phase) in Δ*napM* and wild-type (WT) strains. Deletion of the *napM* gene does not affect BD phase duration (34.67 ± 16.34 min and 36.02 ± 16.41, respectively; $t(1314) = -1.50$ $p = 1.3*10^{-1}$, $n = 658$).

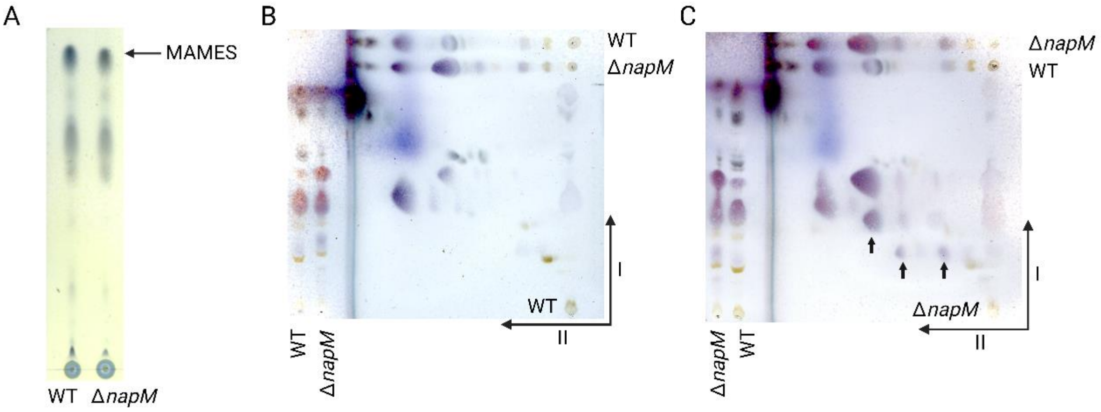


**Figure S4. MAMES TLC (A) and 2D-TLC analysis of the wild-type (WT) (B) and Δ*napM* (C) strains.** Red arrows indicate enrichment in lipids fractions in deletion mutant.

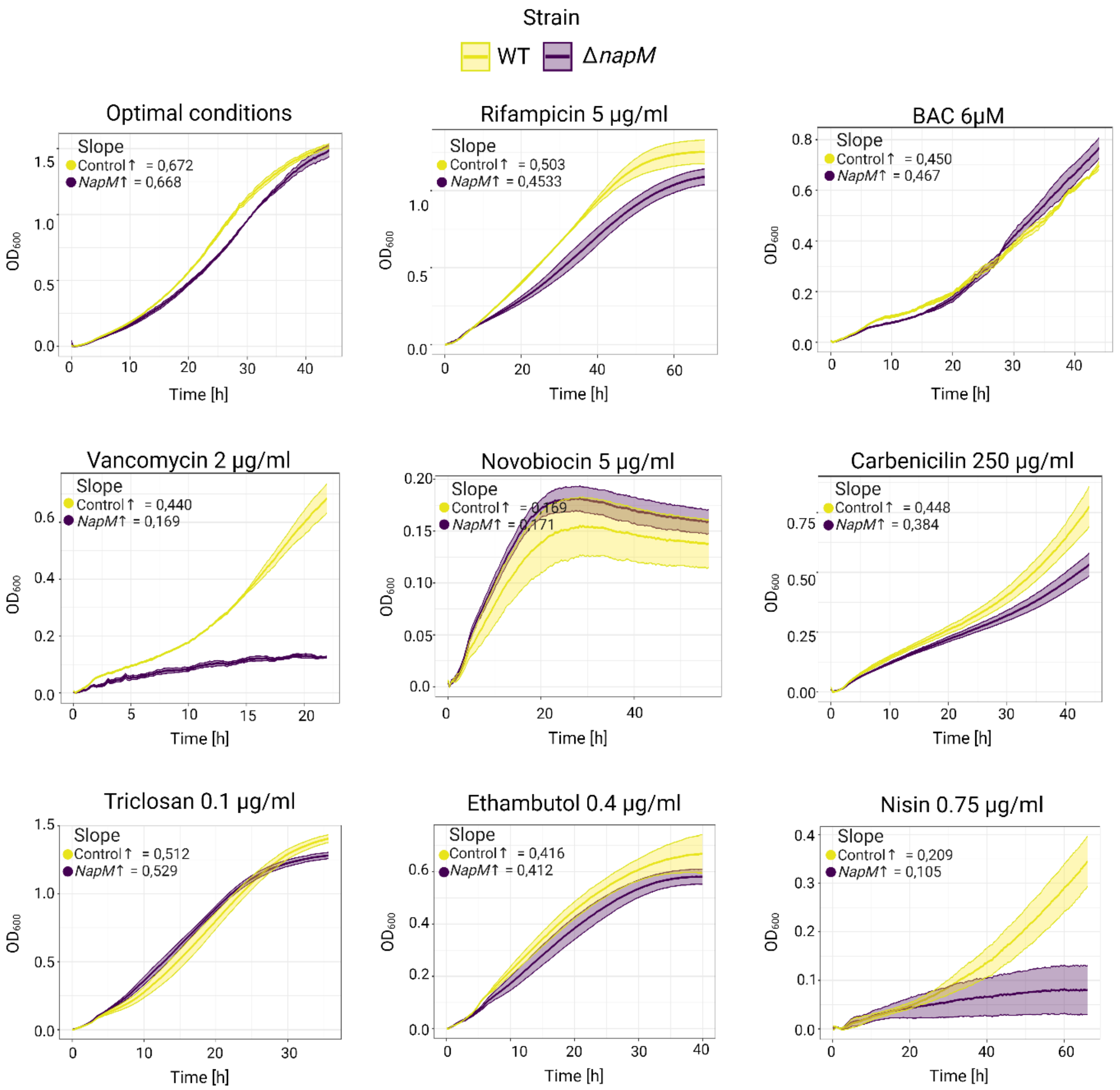


**Figure S5. Analysis of growth rates in *M. smegmatis* cells overproducing NapM under various stress conditions.** Growth curves of the NapM↑ strain compared to the Control↑ strain.

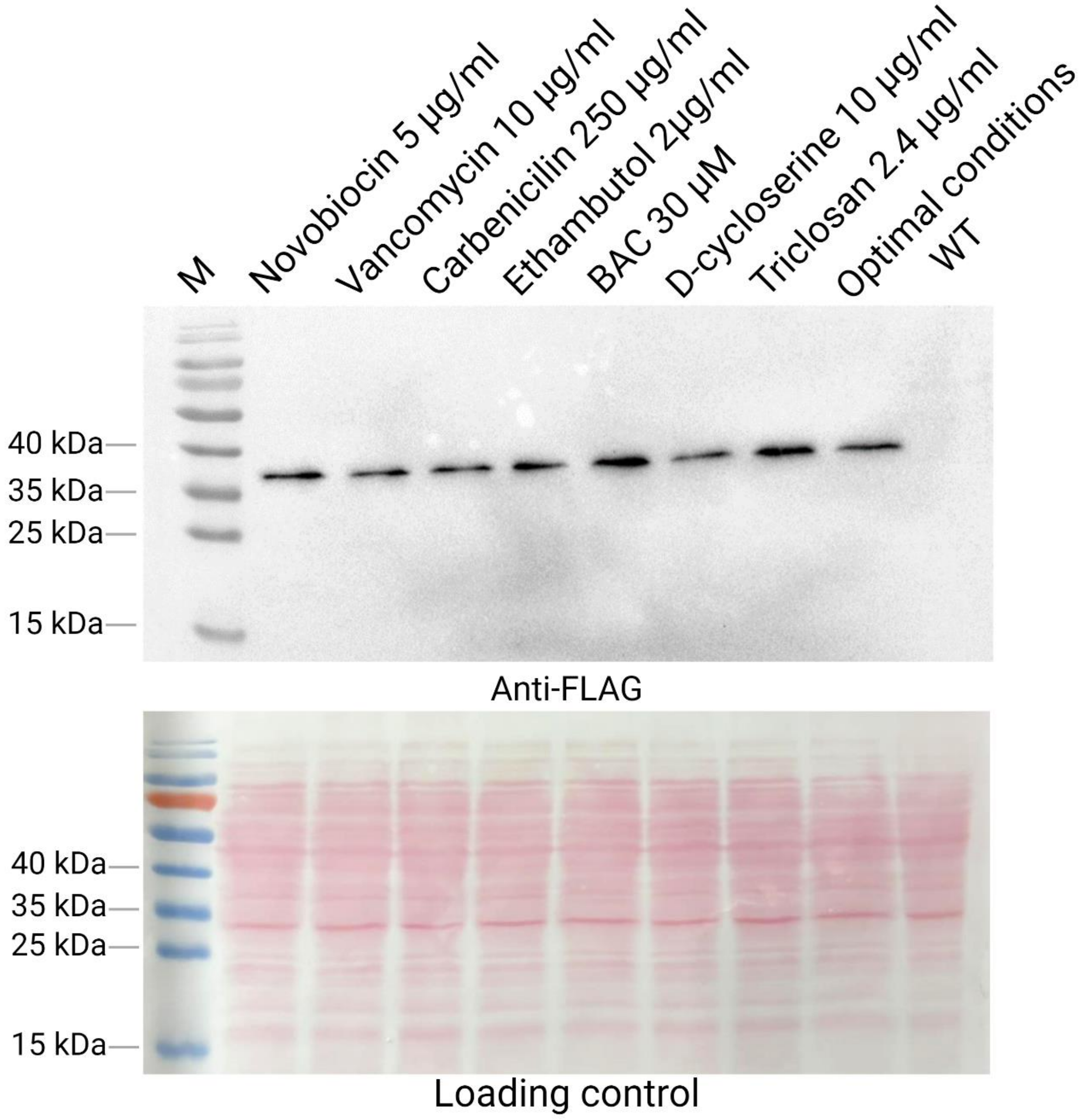


**Figure S6. Western blotting analysis showing upregulation of NapM-FLAGx3 protein upon exposure to stress factors.** Estimated NapM-FLAGx3 protein size is 24 kDa.

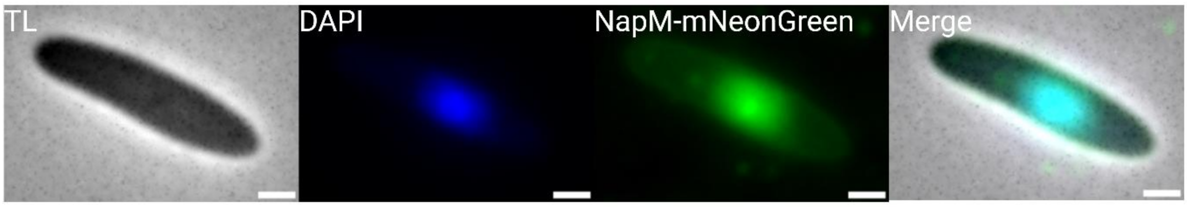


**Figure S7. Colocalization of NapM-mNeonGreen with DAPI-stained nucleoid in *E. coli* cells.** TL- transmitted light. Scale bar, 1 µm

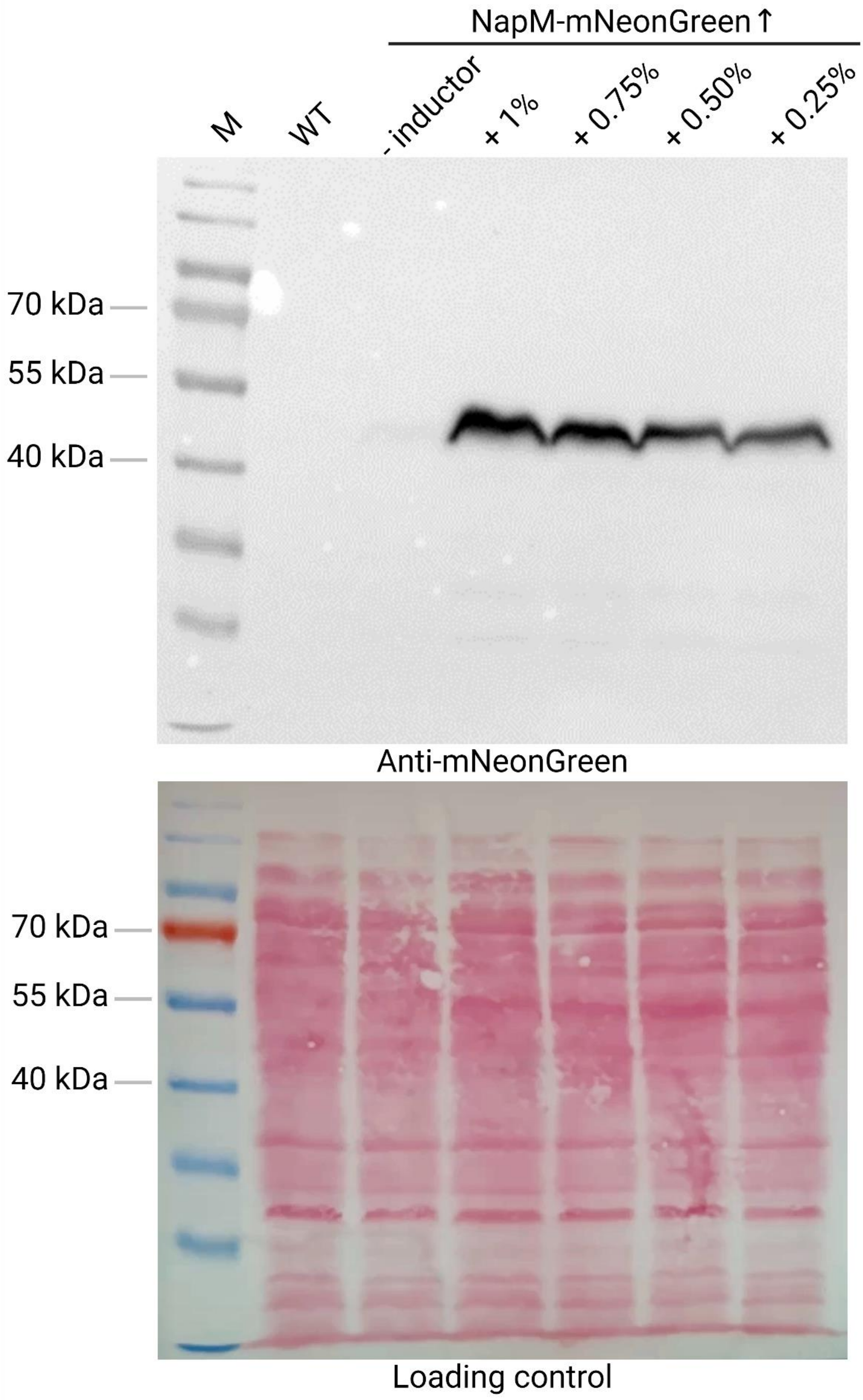


**Figure S8. Western blotting analysis of NapM-mNeonGreen fusion protein level in NapM-mNeonGreen↑ strain in optimal conditions with different acetamide (inducer) concentrations.** Estimated NapM-mNeonGreen protein size is 50 kDa.

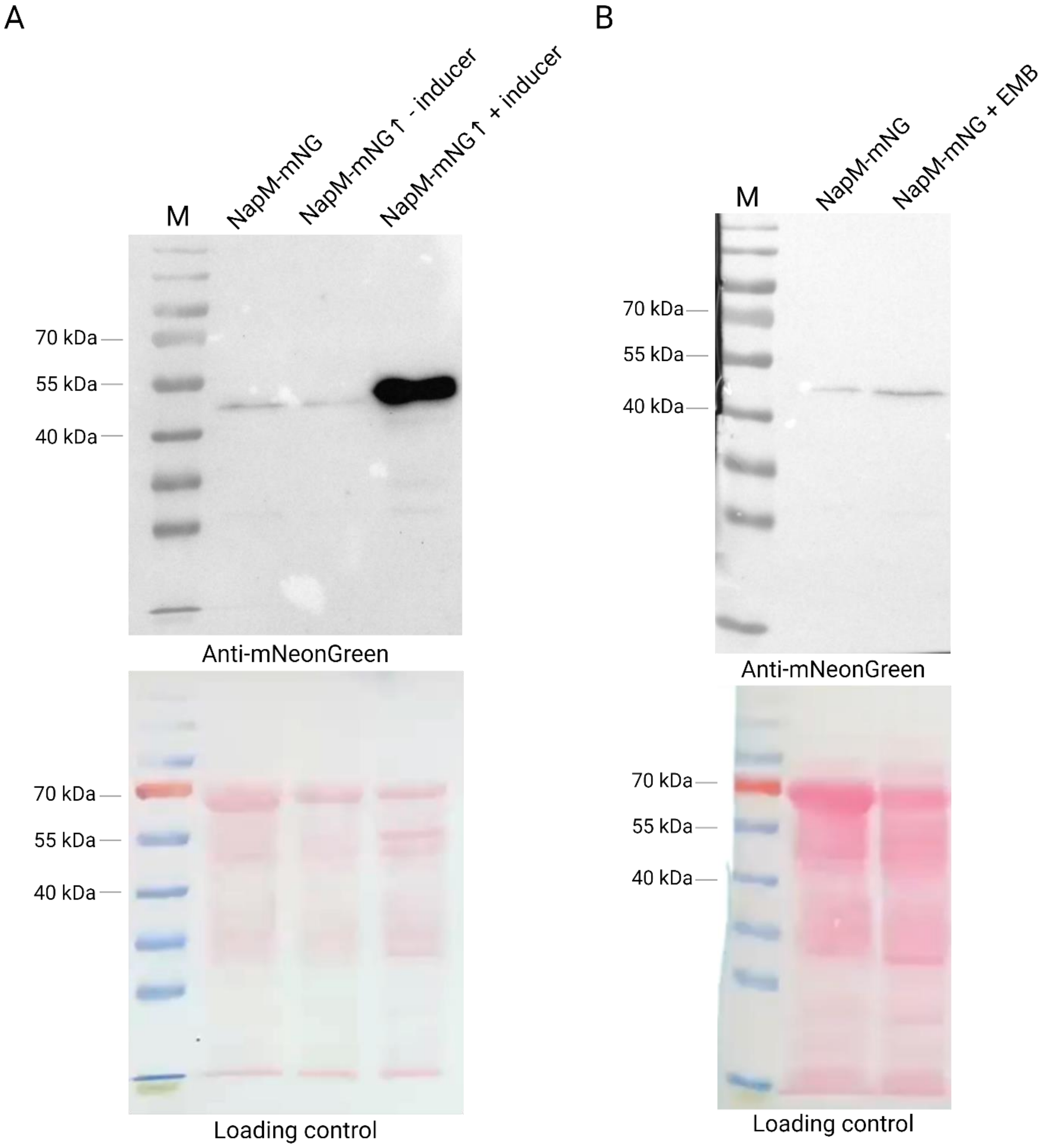


**Figure S9. Western blotting analysis of NapM-mNeonGreen and NapM-mNeonGreen↑ strains.** Estimated protein size is 50 kDa. **A.** Comparison of the levels of NapM-mNeonGreen fusion protein produced in strains expressing *napM-mneongreen* gene under native (NapM-mNeonGreen strain) and inducible promotor (NapM-mNeonGreen↑ strain). **B.** Western blotting analysis of NapM-mNeonGreen level in optimal conditions and after 3 hours exposition to 2 μg/ml ethambutol (EMB).

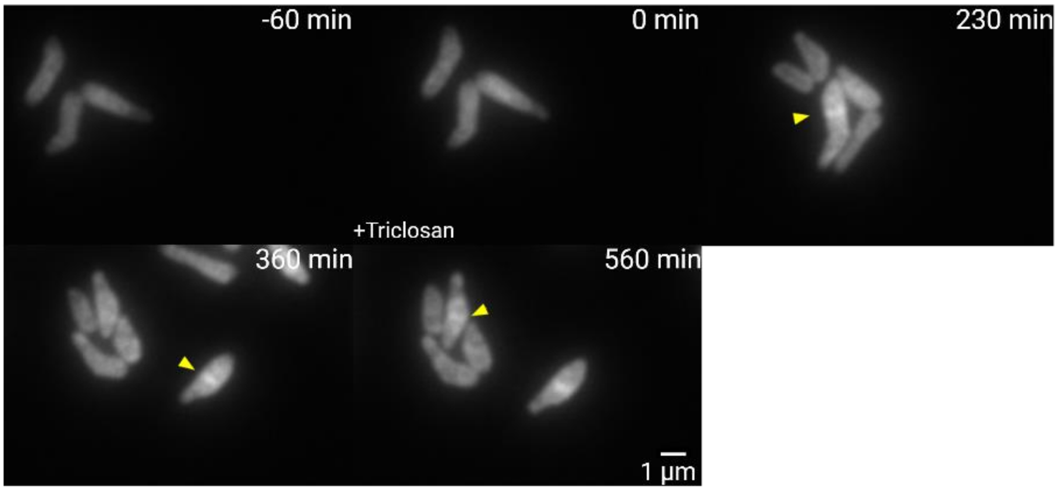


**Figure S10. Time-lapse imaging of NapM-mNeonGreen↑ cells upon 6h Triclosan exposure.** Yellow arrows indicate septal localization of the NapM-mNeonGreen fusion protein.

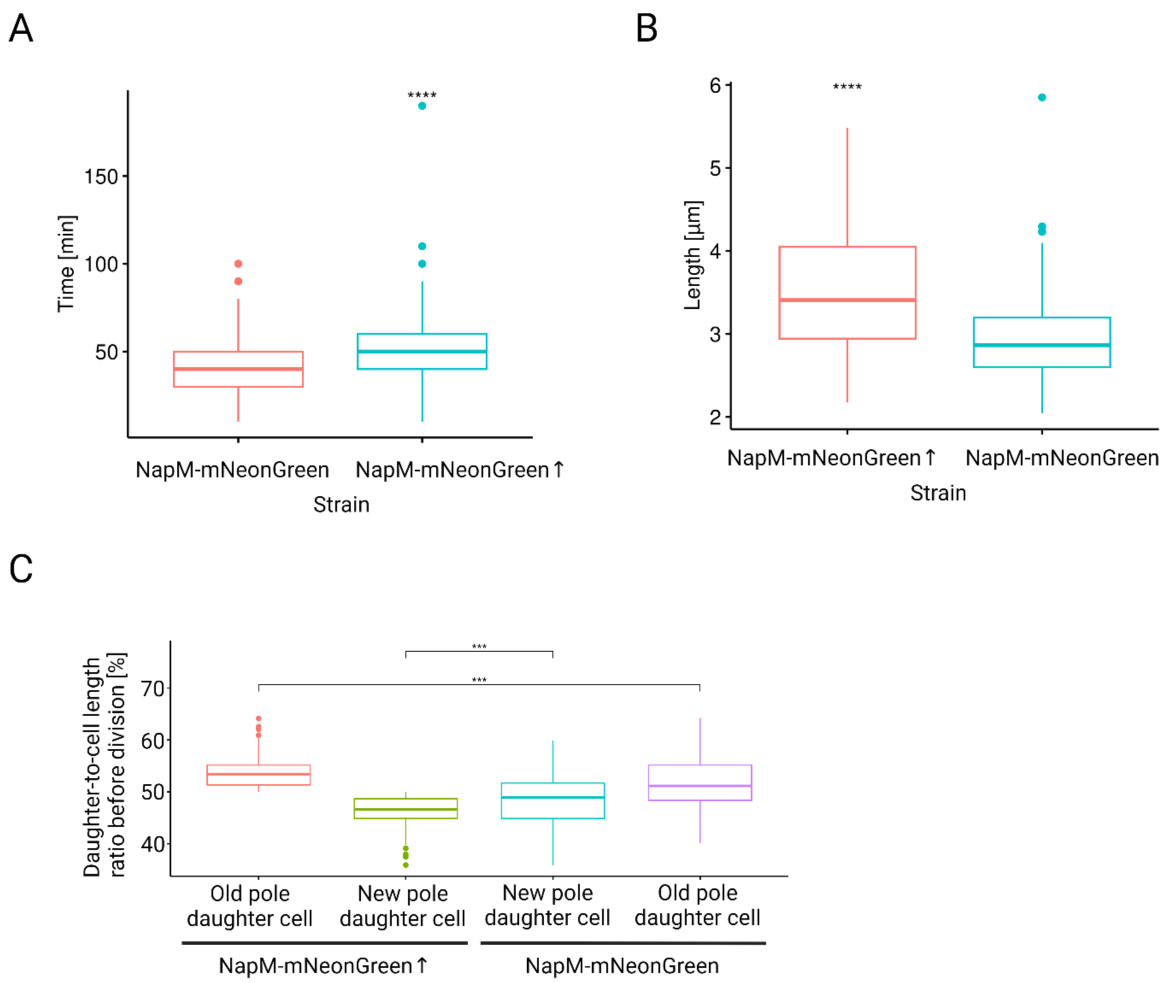


**Figure S11. Analysis of NapM-mNeonGreen localization in *M. smegmatis* cells upon ethambutol exposure**. **A.** Duration of NapM-mNeonGreen signal within the septum area in NapM-mNeonGreen vs. NapM-mNeonGreen↑ cells. **B.** Comparison of NapM-mNeonGreen and NapM-mNeonGreen↑ cells length upon division. **C.** Analysis of daughter cells asymmetry in NapM-mNeonGreen and NapM-mNeonGreen↑ strains. Length of NapM-mNeonGreen daughter cells is more equal (i.e., the division is more symmetrical) than in the case of NapM-mNeonGreen↑ strain.

**Movie 1.** Time-lapse imaging of NapM-mNeonGreen cells using ONIX microfluidics system upon 6h ethambutol (EMB) exposure, followed by antibiotic washing. Yellow arrows indicate septum-like localization of NapM-mNeonGreen. Images were acquired automatically every 10 min. Scale bar, 1 µm.

**Movie 2.** Time-lapse imaging of NapM-mNeonGreen↑ cells using ONIX microfluidics system upon 6h ethambutol (EMB) exposure, followed by antibiotic washing. Yellow arrows indicate septum-like localization of NapM-mNeonGreen. Images were acquired automatically every 10 min. Scale bar, 1 μm.
